# Changes in intra- and interlimb reflexes from hindlimb cutaneous afferents after staggered thoracic lateral hemisections during locomotion in cats

**DOI:** 10.1101/2023.12.15.571869

**Authors:** Stephen Mari, Charly G. Lecomte, Angèle N. Merlet, Johannie Audet, Sirine Yassine, Oussama Eddaoui, Gabriel Genois, Charlène Nadeau, Jonathan Harnie, Ilya A. Rybak, Boris I. Prilutsky, Alain Frigon

**Affiliations:** Department of Pharmacology-Physiology, Faculty of Medicine and Health Sciences, Université de Sherbrooke, Centre de recherche du Centre Hospitalier Universitaire de Sherbrooke, Sherbrooke, QC, Canada; Department of Neurobiology and Anatomy, Drexel University College of Medicine, Philadelphia, PA, United States; School of Biological Sciences, Georgia Institute of Technology, Atlanta, GA, United States

**Author notes:** Corresponding author: Alain Frigon, PhD.

**Keywords:** cutaneous reflexes, interlimb coordination, locomotion, spinal cord injury

## Abstract

When the foot dorsum contacts an obstacle during locomotion, cutaneous afferents signal central circuits to coordinate muscle activity in the four limbs. Spinal cord injury disrupts these interactions, impairing balance and interlimb coordination. We evoked cutaneous reflexes by electrically stimulating left and right superficial peroneal nerves before and after two thoracic lateral hemisections placed on opposite sides of the cord at 9-13 weeks interval in seven adult cats (4 males and 3 females). We recorded reflex responses in ten hindlimb and five forelimb muscles bilaterally. After the first (right T5-T6) and second (left T10-T11) hemisections, coordination of the fore- and hindlimbs was altered and/or became less consistent. After the second hemisection, cats required balance assistance to perform quadrupedal locomotion. Short-latency reflex responses in homonymous and crossed hindlimb muscles largely remained unaffected after staggered hemisections. However, mid- and long-latency homonymous and crossed responses in both hindlimbs occurred less frequently after staggered hemisections. In forelimb muscles, homolateral and diagonal mid- and long-latency response occurrence significantly decreased after the first and second hemisections. In all four limbs, however, when present, short-, mid- and long-latency responses maintained their phase-dependent modulation. We also observed reduced durations of short-latency inhibitory homonymous responses in left hindlimb extensors early after the first hemisection and delayed short-latency responses in the right ipsilesional hindlimb after the first hemisection. Therefore, changes in cutaneous reflex responses correlated with impaired balance/stability and interlimb coordination during locomotion after spinal cord injury. Restoring reflex transmission could be used as a biomarker to facilitate locomotor recovery.

**Key points:**

- Cutaneous afferent inputs coordinate muscle activity in the four limbs during locomotion when the foot dorsum contacts an obstacle.
- Thoracic spinal cord injury disrupts communication between spinal locomotor centers located at cervical and lumbar levels, impairing balance and limb coordination.
- We investigated cutaneous reflexes during quadrupedal locomotion by electrically stimulating the superficial peroneal nerve bilaterally, before and after staggered lateral thoracic hemisections of the spinal cord in cats.
- We showed a loss/reduction of mid- and long-latency responses in all four limbs after staggered hemisections, which correlated with altered coordination of the fore- and hindlimbs and impaired balance.
- Targeting cutaneous reflex pathways projecting to the four limbs could help develop therapeutic approaches aimed at restoring transmission in ascending and descending spinal pathways.

## INTRODUCTION

The control of locomotion involves dynamic interactions between supraspinal structures, spinal circuits and somatosensory feedback to coordinate limb movements and maintain balance/stability [for reviews, see (Rossignol *et al*., 2006; Frigon, 2017; Frigon *et al*., 2021)]. Spinal cord injury (SCI) disrupts these interactions, leading to severe sensorimotor deficits, including impaired balance and limb coordination (Barbeau *et al*., 2002; Edgerton *et al*., 2004; Van Hedel & Dietz, 2010; Rossignol & Frigon, 2011). Although complete SCI abolishes all communication between the brain and spinal sensorimotor circuits located below the lesion, most SCIs are incomplete, and some communication between spinal circuits controlling the arms/forelimbs and legs/hindlimbs remains possible. Somatosensory feedback, proprioceptive and tactile, plays an important role in the recovery of hindlimb locomotion after thoracic SCI (Goldberger, 1977; Muir & Steeves, 1995; Bouyer & Rossignol, 2003*a*, 2003*b*; Smith *et al*., 2006; Takeoka *et al*., 2014). However, its role in limb coordination after incomplete SCI is not clear.

One way to address this issue is to evoke reflexes by stimulating peripheral nerves and recording responses in all four limbs before and after SCI. Studies in cats have shown changes in reflex responses in the hindlimbs following incomplete or complete thoracic SCI (Frigon & Rossignol, 2008; Frigon *et al*., 2009; Gossard *et al*., 2015). However, only a few studies have investigated changes in reflex pathways that send signals between circuits located at cervical and lumbosacral levels that control and coordinate the arms/forelimbs and legs/hindlimbs, respectively, after SCI. One study reported an increased ascending transmission two weeks after a unilateral cervical SCI in rats by electrically stimulating the sciatic nerve and recording responses in a forelimb extensor muscle bilaterally at rest (Côté *et al*., 2012). In humans, interlimb reflexes evoked in arm muscles with electrical stimulation of the tibial nerves at rest, were strengthened following cervical SCI (Calancie *et al*., 2002). We know that interlimb reflexes evoked by stimulating cutaneous nerves of the fore- and hindlimbs are modulated during locomotion in cats and humans (Haridas & Zehr, 2003; Hurteau *et al*., 2018; Merlet *et al*., 2022; Mari *et al*., 2023), likely contributing to limb coordination, but we do not know how they change following SCI and if their loss could explain impairments in limb coordination during locomotion.

In the present study, we electrically stimulated the superficial peroneal nerve (SP) that innervates the foot dorsum, responsible for the stumbling corrective reaction in cats and humans (Forssberg *et al*., 1977; Prochazka *et al*., 1978; Forssberg, 1979; Duysens & Loeb, 1980; Wand *et al*., 1980; Schillings *et al*., 1996; Van Wezel *et al*., 1997; Zehr *et al*., 1997; Quevedo *et al*., 2005*b*, 2005*a*), during treadmill locomotion. In cats, stimulating the SP nerve elicits short- (7-19 ms), mid- (19-34 ms) and long-latency (35-60 ms) responses in all four limbs (Hurteau *et al*., 2018; Merlet *et al*., 2022; Mari *et al*., 2023). Mid- and long-latency responses are thought to involve supraspinal contributions (Fuwa *et al*., 1991; LaBella *et al*., 1992; Frigon & Rossignol, 2008; Hurteau & Frigon, 2018). We stimulated the SP nerve before, after a mid-thoracic lateral hemisection on the right side and then after a second lateral hemisection on the left side a few spinal segments caudal to the first hemisection 9-13 weeks later. This staggered hemisections paradigm disrupts direct ascending and descending spinal pathways that communicate between the brain/cervical cord and lumbosacral circuits, severely impairing fore-hind coordination (Jane *et al*., 1964; Kato *et al*., 1984, 1985; Stelzner & Cullen, 1991; Courtine *et al*., 2008; Van Den Brand *et al*., 2012; Cowley *et al*., 2015; Audet *et al*., 2023). Here, we used this paradigm to probe neural changes in interlimb pathways and to determine their potential role in the recovery of forelimb-hindlimb coordination. We hypothesized that the loss of interlimb reflexes in the forelimbs with SP nerve stimulation parallels the loss of fore-hind coordination, consistent with limited neural communication between cervical and lumbosacral sensorimotor circuits. Our results show that the occurrence of mid- and long-latency reflex responses in all four limbs was considerably reduced after the first and second spinal lesions, particularly in the forelimbs. These changes in cutaneous reflexes correlated with altered coordination between the fore- and hindlimbs during locomotion as well as with a loss in balance.

## MATERIAL AND METHODS

### Ethical approval

All procedures were approved by the Animal Care Committee of the Université de Sherbrooke (Protocol 442-18) in accordance with policies and directives of the Canadian Council on Animal Care. We obtained the current data set from seven adult purpose-bred cats (> 1 year of age at the time of experimentation), 3 females and 4 males, weighing between 3.4 kg and 6.5 kg, purchased from Marshall BioResources. Before and after experiments, cats were housed and fed in a dedicated room within the animal care facility of the Faculty of Medicine and Health Sciences at the Université de Sherbrooke. We followed the ARRIVE guidelines 2.0 for animal studies (Percie Du Sert *et al*., 2020). The investigators understand the ethical principles under which the journal operates and our work complies with this animal ethics checklist. In order to maximize the scientific output of each animal, they were used in other studies to investigate different scientific questions, some of which have been published (Lecomte *et al*., 2022, 2023; Merlet *et al*., 2022; Audet *et al*., 2023; Mari *et al*., 2023).

### General surgical procedures

All surgeries (implantation and spinal lesions) were performed under aseptic conditions with sterilized equipment in an operating room. Prior to surgery, cats were sedated with an intramuscular (i.m.) injection of butorphanol (0.4 mg/kg), acepromazine (0.1 mg/kg), and glycopyrrolate (0.01 mg/kg). We then injected a mixture of diazepam/ketamine (0.05 mg/kg, i.m.) five minutes later for induction. We shaved the animal’s fur (back, stomach, fore- and hindlimbs) and cleaned the skin with chlorhexidine soap. Cats were anesthetized with isoflurane (1.5-3%) and O2 delivered with a mask and then with a flexible endotracheal tube. Anesthesia was maintained during surgery by adjusting isoflurane concentration as needed and by monitoring cardiac and respiratory rates. Body temperature was maintained constant (37 ± 0.5°C) using a water-filled heating pad placed under the animal, an infrared lamp placed ∼50 cm over it and a continuous infusion of lactated Ringers solution (3 ml/kg/h) through a catheter placed in a cephalic vein. At the end of surgery, we injected subcutaneously an antibiotic (cefovecin, 8 mg/kg) and a fast-acting analgesic (buprenorphine, 0.01 mg/kg). We also taped a fentanyl (25 µg/h) patch to the back of the animal 2-3 cm rostral to the base of the tail for prolonged analgesia, which we removed 4-5 days later. After surgery, cats were placed in an incubator and closely monitored until they regained consciousness. We administered another dose of buprenorphine ∼7 hours after surgery. At the end of experiments, cats were administered a lethal dose of pentobarbital (120 mg/kg) under general anesthesia through the left or right cephalic vein and spinal cords were harvested for histological analysis (Audet *et al*., 2023; Lecomte *et al*., 2023).

### Staggered hemisections

After collecting data in the intact state and using the same general surgical procedures described above, we performed a lateral hemisection between the 5^th^ and 6^th^ thoracic vertebrae (T5-T6) on the right side of the spinal cord. An incision of the skin over T5-T6 was made and after carefully setting aside muscle and connective tissue, a small laminectomy of the corresponding dorsal bone was performed. Lidocaine (xylocaine, 2%) was applied topically followed by 2-3 intraspinal injections on the right side of the cord. We then sectioned the spinal cord laterally from the midline to the right using surgical scissors. We placed hemostatic material (Spongostan) within the gap before sewing back muscles and skin in anatomical layers. In the days following hemisection, voluntary bodily functions were carefully monitored. The bladder and large intestine were manually expressed if needed. Once data were collected following the first hemisection (9-13 weeks), we performed a second lateral hemisection between the 10^th^ and 11^th^ thoracic vertebrae (T10-T11) on the left side of the spinal cord using the same surgical procedures and post-operative care described above.

### Electromyography and nerve stimulation

To record the electrical activity of muscles (EMG, electromyography), we directed pairs of Teflon-insulated multistrain fine wires (AS633; Cooner Wire) subcutaneously from two head-mounted 34-pin connectors (Omnetics Connector). Two wires, stripped of 1–2 mm of insulation, were sewn into the belly of selected forelimb/hindlimb muscles for bipolar recordings. The head-mounted connectors were fixed to the skull using dental acrylic and four to six screws. We verified electrode placement during surgery by stimulating each muscle through the appropriate head connector channel to assess the biomechanically desired muscle contraction. During experiments, EMG signals were pre-amplified (×10, custom-made system), bandpass filtered (30–1,000 Hz) and amplified (100–5,000×) using a 16-channel amplifier (model 3500; AM Systems). EMG data were digitized (5,000 Hz) with a National Instruments card (NI 6032E), acquired with custom-made acquisition software and stored on computer. Five forelimb muscles were implanted bilaterally: biceps brachii (BB, elbow and shoulder flexor), the long head of the triceps brachii (TRI, elbow and shoulder extensor), latissimus dorsi (LD, shoulder retractor), extensor carpi ulnaris (ECU, wrist dorsiflexor) and flexor carpi ulnaris (FCU, wrist plantarflexor). Ten hindlimb muscles were implanted bilaterally: anterior sartorius (SRT, hip flexor and knee extensor), semitendinosus (ST, knee flexor and hip extensor), vastus lateralis (VL, knee extensor), iliopsoas (IP, hip flexor), biceps femoris posterior (BFP, hip extensor and knee flexor), biceps femoris anterior (BFA, hip extensor), lateral gastrocnemius (LG, ankle plantarflexor and knee flexor), soleus (SOL, ankle plantarflexor), medial gastrocnemius (MG, ankle plantarflexor and knee flexor), and tibialis anterior (TA, ankle dorsiflexor).

For bipolar nerve stimulation, pairs of Teflon-insulated multistrain fine wires (AS633; Cooner Wire) were passed through a silicon tubing. A horizontal slit was made in the tubing and wires within the tubing were stripped of their insulation. The ends protruding through the cuff were knotted to hold the wires in place and glued. The ends of the wires away from the cuff were inserted into four-pin connectors (Hirose or Samtec) and fixed to the skull using dental acrylic. Cuff electrodes were directed subcutaneously from head-mounted connectors to the left and right SP nerves at the ankle which are purely cutaneous at these levels (Bernard *et al*., 2007).

### Experimental design

We collected EMG and kinematic data before and at different time points after staggered hemisections during quadrupedal locomotion at the cat’s preferred treadmill speed (0.3-0.5 m/s). Cats KA and KI stepped at 0.3 and 0.5 m/s, respectively, while the other five cats stepped at 0.4 m/s. The treadmill consisted of two independently controlled belts 130 cm long and 30 cm wide (Bertec) with a Plexiglas separator (130 cm long, 7 cm high, and 1.3 cm wide) placed between the two belts to prevent limbs impeding each other. In the intact, preoperative state, cats were trained for 2-3 weeks in a progressive manner, first for a few steps and then for several consecutive minutes, using food and affection as rewards. Once cats could perform 3-4 consecutive minutes, we started the experiments. During experiments, we delivered trains of electrical stimuli consisting of three 0.2 ms pulses at 300 Hz using a Grass S88 stimulator. At the start of the experiment, we determined the motor threshold, defined as the minimal intensity that elicited a small motor response in an ipsilateral flexor muscle (e.g., ST or TA) during the swing phase. We then set stimulation intensity at 1.2 times the motor threshold. A locomotor trial lasted 4-5 min and consisted of ∼60 stimuli delivered pseudo-randomly every 2–4 cycles at four different locomotor phases, based on the onset of an extensor burst.

Stimulation deliveries corresponded approximately to mid-stance, the stance-to-swing transition, mid-swing and the swing-to-stance transition of the stimulated limb, which were established at the start of the experiment based on the extensor burst onset. We characterized responses in muscles of the stimulated limb (homonymous), the opposite limb of the same girdle (crossed), the limb on the same side (homolateral) and the diagonal limb (diagonal). **Figure 1A** describes the timeline of data collection in all cats at two time points after the first hemisection (**H1T1** and **H1T2**) and at 1-2 time points after the second hemisection (**H2T1** in 2 cats and **H2T2** in 6 cats). Some cats only have one time point after the second hemisection because they took longer to recover quadrupedal locomotion. No data were collected for cat KI after the second hemisection due to technical issues with the implants.

**Figure 1.**
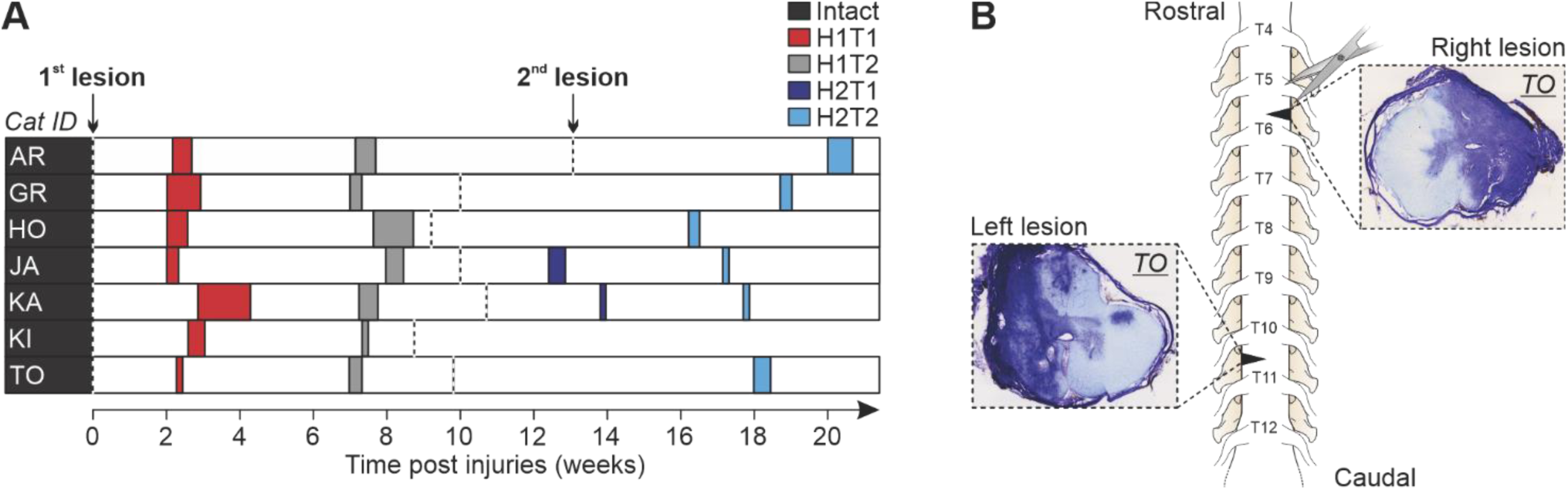
Experimental timeline and staggered hemisections paradigm. (**A**) Chronology showing the first (T1) and second (T2) experimental time points after the first (H1) and second (H2) hemisections in all cats. (B) Schematic representation of the staggered hemisections, with the first and second hemisections at right (T5–T6) and left (T10–T11) thoracic levels, respectively. Spinal lesions (cat TO) are highlighted with Cresyl violet staining.

### Histology

After confirming euthanasia (i.e., no cardiac and respiratory functions), we harvested an approximately 2 cm long section of the spinal cord centered on the lesions. Segments of the dissected spinal cord were then placed in a 25 ml 4% paraformaldehyde solution (PFA; 0.1 M PBS, 4°C). After 5 days, we placed the spinal cord in a new PBS (0.2 M) solution containing 30% sucrose for 72 h at 4°C, then froze it in isopentane at - 50°C for cryoprotection. The spinal cord was then sliced in 50 µm coronal sections using a cryostat (Leica CM1860) and mounted on gelatinized-coated slides. The slides were dried overnight and then stained with a 1% cresyl violet acetate solution for 12 min. We washed the slides for 3 min in distilled water before being dehydrated in successive baths of ethanol (50%, 70% and 100%, 5 min each) and transferring them in xylene for 5 min. Dibutylphthalate polystyrene xylene was next used to mount and dry the spinal cord slides before being scanned by a Nanozoomer. We then performed qualitative and quantitative analyses to estimate lesion extent using ImageJ by selecting the slide with the greatest identifiable damaged area. Using the scarring tissue stained with cresyl violet acetate, we estimated lesion extent by dividing the lesion area by the total area of the selected slice and expressed it as percentage. **Figure 1B** shows a schematic of the staggered hemisections with histological cross-section images for the first and second lesion in a representative cat.

### Kinematic acquisition and analysis

During experiments, two cameras (Basler AcA640-100gm) captured videos from the left and right sides of the animals (60 frames per second; 640 × 480 pixels spatial resolution). A custom-made LabVIEW program acquired the images and synchronized the cameras with EMG data. We analyzed kinematic data from videos off-line with a deep learning approach (DeepLabCutTM; Mathis et al., 2018), which we recently validated in our cat model (Lecomte *et al*., 2021). We determined contact and liftoff of the four limbs by visual inspection. Paw contact was defined as the first frame where the paw made visible contact with the treadmill surface. Paw liftoff was defined as the frame with the most caudal displacement of the toe. We measured cycle duration as the interval of time from successive paw contacts of the same limb. Stance duration corresponded to the interval of time from contact to liftoff of the same limb, while swing duration was measured as cycle duration minus stance duration. We quantified temporal fore-hind coordination by measuring the phase interval between contact of the right hindlimb and right forelimb divided by cycle duration of the right hindlimb (English, 1979; English & Lennard, 1982; Orsal *et al*., 1990; Frigon *et al*., 2014; Thibaudier & Frigon, 2014; Thibaudier *et al*., 2017; Audet *et al*., 2022, 2023). After the first and second hemisections, cats often performed 2:1 fore-hind patterns (i.e., two right forelimb cycles for one right hindlimb cycle) and we separated the first and second forelimb cycles when this occurred. Phase intervals were then multiplied by 360 and expressed in degrees to illustrate their periodic nature and possible distributions (English & Lennard, 1982). Functionally, phase interval values indicate when the right forelimb made contact relative to right hindlimb contact.

### Reflex analysis

We describe the reflex analysis in several of our publications (Hurteau *et al*., 2017, 2018; Hurteau & Frigon, 2018; Merlet *et al*., 2020, 2021; Mari *et al*., 2023). The step-by-step procedure for quantifying reflex responses is illustrated for the left soleus with stimulation of the left SP nerve (**Fig. 2**). For all locomotor sessions, EMG signals were low-pass filtered (250 Hz) to facilitate the visualization of the EMG activity envelope. We first defined locomotor cycles from successive burst onsets of the left sartorius and separated them as stimulated (i.e., cycles with stimulation) or control (i.e., cycles without stimulation) cycles. Sections where the cat stepped irregularly were removed from analysis based on EMG and video data. Stimulated cycles were then sorted and divided into 4 subphases based on stance onset of the stimulated limb: swing-to-stance, mid-stance, stance-to-swing and mid-swing. Control (C̅) cycles were averaged and rectified to provide a baseline locomotor EMG, an indication of the excitability level of the motor pool at stimulation. We averaged the stimulated (S̅) cycles and time normalized C̅ to S̅ cycle durations and superimposed them. To determine response onsets and offsets, defined as prominent positive or negative deflections away from C̅, we set windows using previous studies as guidelines (Duysens & Stein, 1978; Duysens & Loeb, 1980; Pratt *et al*., 1991; Loeb, 1993; Hurteau *et al*., 2017, 2018; Hurteau & Frigon, 2018; Mari *et al*., 2023) with 97.5% confidence intervals. We measured the latencies of responses from stimulation onset to response onset. We also measured response durations as the time interval between response onset and offset. We termed short-latency (7–18 ms; **SLR**) excitatory and inhibitory responses as P1 and N1 responses, respectively, based on the terminology introduced by (Duysens & Loeb, 1980). Responses in the crossed, homolateral, and diagonal limbs that had an onset ≤18 ms were classified as P1 or N1, as the minimal latency for spino-bulbo-spinal reflexes in the cat is 18 ms (Shimamura & Livingston, 1962). Mid-latency (19–34 ms; **MLR**) excitatory and inhibitory responses were termed P2 and N2, respectively. Long-latency (35–60 ms; **LLR**) excitatory and inhibitory responses were termed P3 and N3, respectively. The EMG of reflex responses S̅ was then integrated and subtracted from the integrated C̅ in the same time window to provide a net reflex value. This net reflex value was then divided by the integrated C̅ value to evaluate reflex responses. This division helps identify if changes in reflex responses across the cycle are independent of changes in C̅ activity (Matthews, 1986; Frigon & Rossignol, 2007, 2008, 2009; Hurteau *et al*., 2017, 2018; Hurteau & Frigon, 2018; Mari *et al*., 2023).

**Figure 2.**
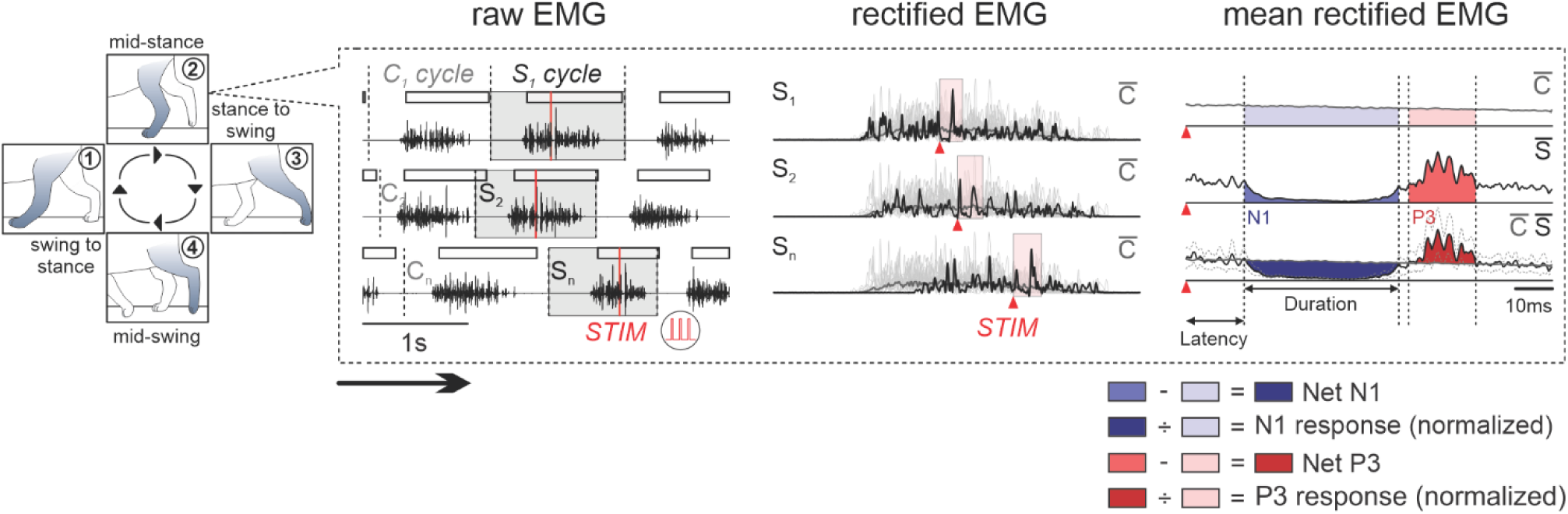
Reflex methodology. Electromyographic (EMG) bursts in a single intact cat (AR) are shown for the left soleus during locomotion with stimulation of the left superficial peroneal nerve at mid-stance. From the *<u>raw EMG</u>* waveforms, we tagged cycles as stimulated (S) when a stimulus fell within the cycle or control if it was not preceded by a stimulated cycle. *<u>Rectified EMG.</u>* We averaged and rectified control cycles (dark gray trace; C̅) aligned to ipsilateral sartorius onset. Each stimulated cycle (black trace) is rectified then time-normalized to match duration from C̅. *<u>Mean rectified EMG.</u>* Stimulated cycles (black trace; S̅) were then averaged and superimposed on the baseline provided from C̅ (dark gray trace). This allowed us to determine positive (in red) and negative (in blue) responses. Onsets and offsets of responses, defined as a prominent positive or negative deflection away from the baseline C̅, were determined visually using confidence intervals at 97.5% (gray dashed traces). The baseline C̅ occurring in the same time window as the response was subtracted from the response in the stimulated cycles S̅ to provide a net reflex value. This value was then divided by the baseline C̅ occurring in the same time window giving N1 and P3 normalized amplitudes for the left soleus.

### Statistical analysis

We performed statistical tests with IBM SPSS Statistics V26 (IBM Corp., Armonk, NY, USA). We quantified reflex responses in five forelimb muscles (BB, ECU, FCU, LD, and TRI) and in ten hindlimb muscles (BFA, BFP, IP, LG, MG, SRT, SOL, ST, TA, and VL) when stimulation was delivered to the left or right SP. To evaluate whether homonymous, crossed, homolateral and diagonal responses were modulated by phase, we performed a one factor (phase) ANOVA on all responses (P1, P2, P3, N1, N2 and N3) in each cat and state/time point. Because we have several responses within a given phase, we considered all responses during a locomotor session as a population. In our statistical analysis, we used mixed models to deal with incomplete data sets. For instance, reflex responses are sometimes absent after spinal lesions.

Response occurrence probabilities, defined as the fraction of evoked responses obtained out of all cats for pooled SLR, MLR and LLR from the different states/time points were compared using a generalized linear mixed model (GLMM) with a binomial distribution and a logit link (mixed logistic regression) in all four limbs. Dependent variables, such as response latencies and durations of homonymous SLR, MLR and LLR, were analyzed using linear mixed model (LMM). The GLMM and LMM analyses were performed using state/time point as a fixed factor. We incorporated random intercepts at two distinct levels to consider the hierarchical relationships present in our dataset. A random intercept on individual cats at the upper level to capture variability across cats. A random intercept on muscle nested within cat at a lower level, acknowledging that the same muscle response data were repeatedly measured within each cat to help us account for any correlation or non-independence of observations within the same cat-muscle pair. For phase intervals at each state/time point, we performed a circular analysis with a MATLAB toolbox for circular statistics (Berens, 2009). We calculated the resultant vector length (r) to quantify the circular spread of values around the mean. A value close to 1 indicates that the data sample is concentrated around the mean direction, whereas a value of 0 indicates a uniform distribution. Rayleigh’s test was then performed to detect unimodal deviation from uniformity from the resultant vector length. Statistical significance for all tests was set at p < 0.05.

## RESULTS

### Recovery of quadrupedal locomotion after staggered hemisections and extent of spinal lesions

Histological analysis shows lesion extent estimations for individual cats after the first and second spinal lesions (**Fig. 3**). They ranged from 40.7% to 66.4% (49.2 ± 8.9%) and 33.5% to 53.7% (46.0 ± 7.6%) for the first and second hemisections, respectively. After the first hemisection, all seven cats regained quadrupedal locomotion on the treadmill within one to two weeks. They were able to perform reflex sessions for several consecutive minutes at the first (H1T1) and second (H1T2) time points. After the second hemisection, of the six cats tested, all recovered quadrupedal locomotion within two to five weeks. However, they required mediolateral balance assistance during reflex sessions, provided by an experimenter that held the tail of the animal but without supporting its weight. As stated in the Methods, some cats only have one time point after the second hemisection because they took longer to recover quadrupedal locomotion. Only two cats, KA and JA, participated in reflex sessions at the first time point (H2T1), i.e., approximately two weeks after the second hemisection. All six cats performed reflex sessions eight weeks later at the second time point (H2T2).

**Figure 3.**
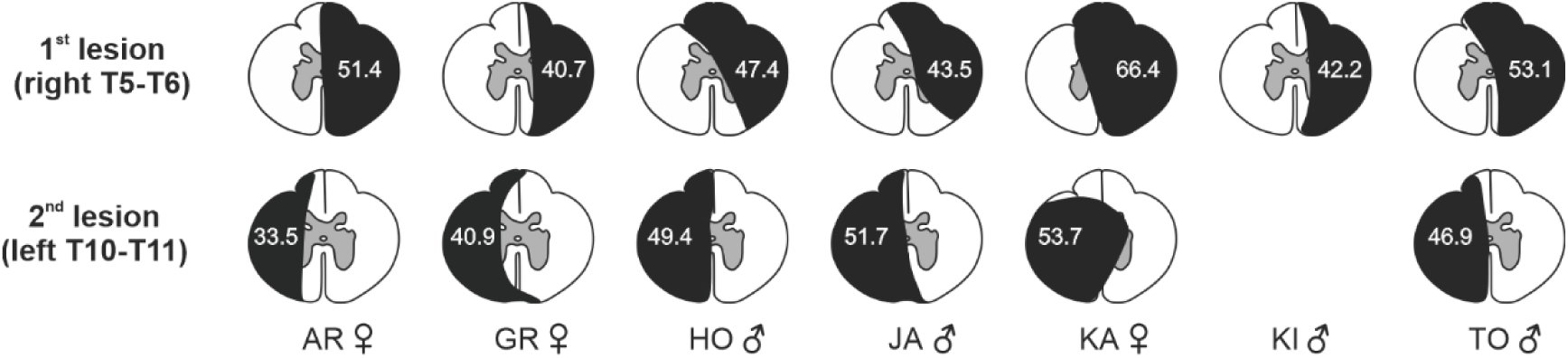
Estimation of the extent of the first and second lesions for individual cats. The black area represents the estimation as a percentage of total cross-sectional area. Note that we only performed one lesion in cat KI.

### Coordination of the fore- and hindlimbs before and after staggered hemisections

After staggered hemisections, the locomotor pattern, including cycle, stance and swing durations, as well as EMG activities changed, as shown for a single cat for illustrative purposes (**Fig. 4**). These changes are described in Audet et al. (2023). In the present study, we focused on changes in the coordination between the fore- and hindlimbs and on reflexes evoked in the four limbs.

**Figure 4.**
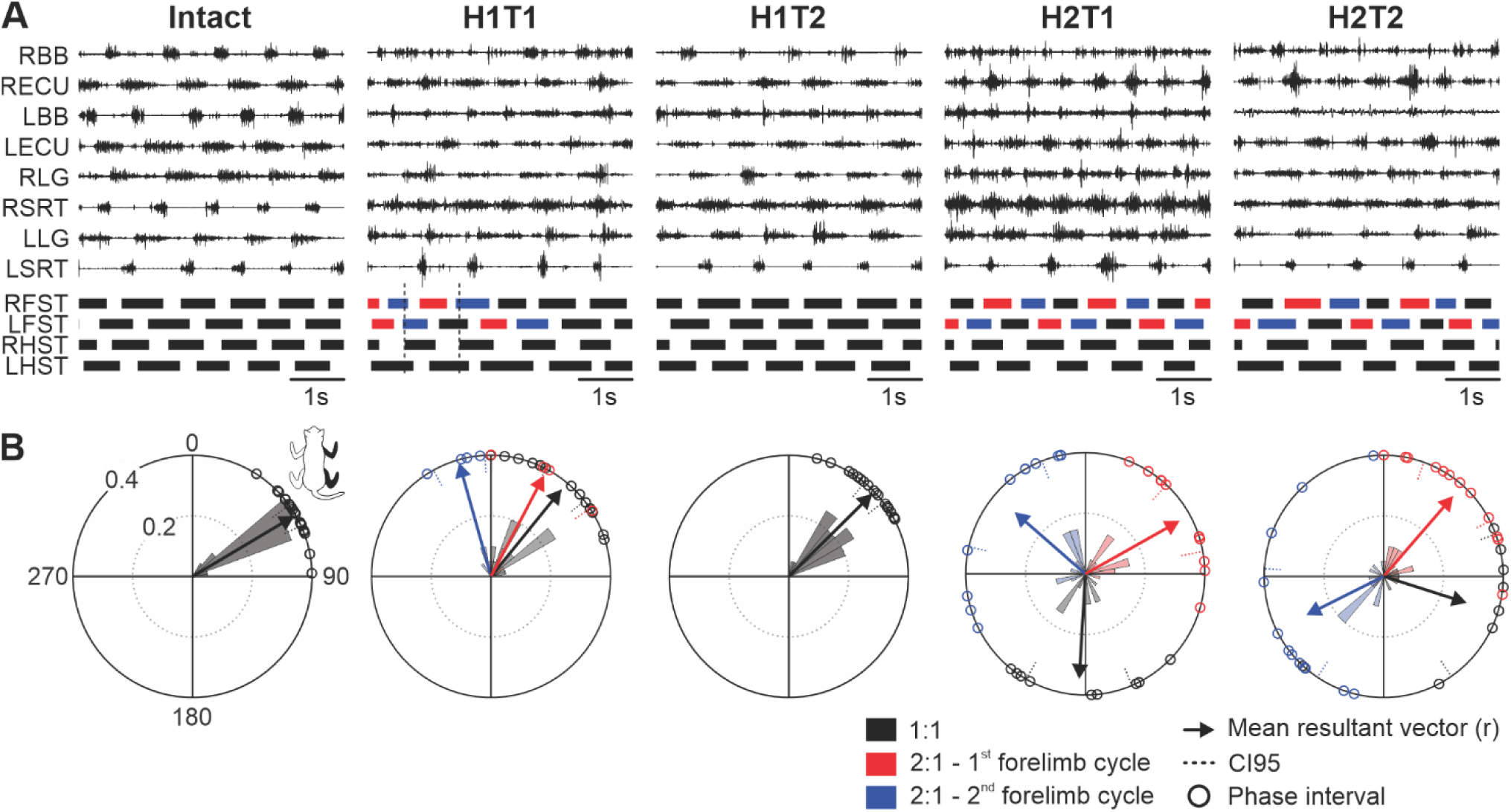
Modulation of the locomotor pattern and fore-hind coordination in a single cat before and after staggered hemisections. (**A**) The figure shows EMG from selected muscles along with the stance phases (thick horizontal bars) of the four limbs in the intact state, and the first (T1) and second (T2) time points after the first (H1) and second (H2) hemisections. Red and blue stance phases represent the first and second forelimb cycles, respectively, during 2:1 fore-hind coordination. The vertical scale is the same for a given muscle in all five panels. (**B**) Circular plots showing phase intervals of right homolateral limbs from 20 control cycles are expressed in degrees around the circumference. Circular histograms plotted in radii indicate the proportion (ratio) of phase interval values to fall within a 10° range. Arrows starting from the center to the circumference indicate mean resultant vectors, i.e., the mean direction where phase intervals are concentrated. The longer their length (r), the greater the concentration of phase interval values around their mean direction. Cycles with 1:1 fore-hind coordination (black) were separated from those with 2:1 coordination. With 2:1 coordination, the first forelimb cycle (red) was separated from the second forelimb cycle (blue). BB, biceps brachii; CI95, 95% confidence intervals; ECU, extensor carpi ulnaris; LFST, left forelimb stance; LG, lateral gastrocnemius; LHST, left hindlimb stance; RFST, right forelimb stance; RHST, right hindlimb stance; SRT, anterior sartorius. Data are from cat KA.

In the intact state, all cats performed 1:1 fore-hind patterns (i.e. one forelimb cycle during a hindlimb cycle), as illustrated for a single cat in **Figure 4A**. In contrast, following the first and/or second hemisection, 2:1 fore-hind patterns could occur, where the right forelimb performed two cycles within a right hindlimb cycle. The proportion of 1:1 and 2:1 fore-hind patterns varied across states/time points and between cats, as summarized in **Table 1**. To determine if 1:1 and 2:1 fore-hind patterns remained coordinated, we measured right homolateral phasing on twenty control cycles (i.e., without stimulation) and performed circular statistics. We separated 1:1 and 2:1 patterns, as well as the first and second right forelimb cycles with 2:1 patterns. As we can see for a representative cat, phase intervals clustered between 30 and 90° (mean of 60 ± 13°) in the intact state, with only 1:1 patterns (**Fig 4B**). At H1T1, 1:1 and 2:1 patterns were present and phase intervals had a mean of 39° (± 23°) for 1:1 patterns and 28° (± 23°) and 344° (± 12°) for the first and second forelimb cycles for 2:1 patterns, respectively. At H1T2, only 1:1 fore-hind patterns were present with a mean around 45° (± 14°). After the second hemisection, 1:1 and 2:1 patterns were again present, with mean phase interval values of 183° (± 31°) for 1:1 patterns and 61° (± 27°) and 309° (± 41°) for the first and second forelimb cycles for 2:1 patterns at H2T1 and 115° (± 51°) for 1:1 patterns and 42° (± 29°) and 250° (± 49°) for the first and second forelimb cycles for 2:1 patterns at H2T2. Thus, fore-hind coordination is considerably altered after the first and second hemisections.

**Table 1.** Proportion of 1:1 and 2:1 fore-hind patterns before and after staggered hemisections. The table shows the proportion of control cycles with 1:1 and 2:1 (first and second right forelimb cycles) fore-hind patterns in individual cats. Proportions and r values (resultant vector length) are represented in the intact state, and after the first (H1) and second (H2) hemisections at 1-2 time points (T1 or T2). An r value close to 1 indicates concentration around the mean direction, whereas a value of 0 indicates uniform distribution. Asterisks indicate a common mean direction of right homolateral phasing (significant Rayleigh’s test, p < 0.05*, p < 0.01** and p < 0.001***).

| Cat ID | State | 1:1 |  | 2:1 |  |  |
| --- | --- | --- | --- | --- | --- | --- |
|  |  | Proportion | *r* | Proportion | *r* (1<sup>st</sup> cycle) | *r* (2<sup>nd</sup> cycle) |
| AR | Intact | 20/20 (100%) | 0.95*** | - | - | - |
|  | H1T1 | 5/20 (25%) | 0.77* | 15/20 (75%) | 0.95*** | 0.81*** |
|  | H1T2 | - | - | 20/20 (100%) | 0.89*** | 0.81*** |
|  | H2T2 | 6/20 (30%) | 0.84** | 14/20 (70%) | 0.92*** | 0.81*** |
| GR | Intact | 20/20(100%) | 0.98*** | - | - | - |
|  | H1T1 | 18/20 (90%) | 0.81*** | 2/20 (10%) | 1.00 | 0.99 |
|  | H1T2 | 16/20 (80%) | 0.77*** | 4/20 (20%) | 0.99** | 0.96* |
|  | H2T2 | 12/20 (60%) | 0.63** | 8/20 (40%) | 0.89*** | 0.62* |
| HO | Intact | 20/20(100%) | 0.95*** | - | - | - |
|  | H1T1 | 16/20 (80%) | 0.42 | 4/20 (20%) | 0.95* | 0.93* |
|  | H1T2 | 20/20 (100%) | 0.61*** | - | - | - |
|  | H2T2 | 10/20 (50%) | 0.75** | 10/20 (50%) | 0.96*** | 0.75** |
| JA | Intact | 20/20 (100%) | 0.95*** | - | - | - |
|  | H1T1 | 13/20 (65%) | 0.52* | 7/20 (35%) | 0.97*** | 0.89*** |
|  | H1T2 | 14/20 (70%) | 0.33 | 6/20 (30%) | 0.95** | 0.89** |
|  | H2T1 | 3/20 (15%) | 0.47 | 17/20 (85%) | 0.85*** | 0.70*** |
|  | H2T2 | 7/20 (35%) | 0.40 | 13/20 (65%) | 0.93*** | 0.70*** |
| KA | Intact | 20/20 (100%) | 0.98*** | - | - | - |
|  | H1T1 | 16/20 (80%) | 0.93*** | 4/20 (20%) | 0.94* | 0.98** |
|  | H1T2 | 20/20 (100%) | 0.97*** | - | - | - |
|  | H2T1 | 9/20 (45%) | 0.87*** | 11/20 (55%) | 0.90*** | 0.78*** |
|  | H2T2 | 8/20 (40%) | 0.73* | 12/20 (60%) | 0.88*** | 0.72** |
| KI | Intact | 20/20 (100%) | 0.97*** | - | - | - |
|  | H1T1 | 20/20 (100%) | 0.96*** | - | - | - |
|  | H1T2 | 20/20 (100%) | 0.97*** | - | - | - |
| TO | Intact | 20/20 (100%) | 0.73*** | - | - | - |
|  | H1T1 | 7/20 (35%) | 0.33 | 13/20 (65%) | 0.81*** | 0.64*** |
|  | H1T2 | 5/20 (25%) | 0.84* | 15/20 (75%) | 0.76*** | 0.77*** |
|  | H2T2 | 4/20 (20%) | 0.15 | 16/20 (80%) | 0.81*** | 0.68*** |

To assess the step-by-step consistency of right homolateral phasing, we performed Rayleigh’s test on the resultant vector length (r) of phase interval values (**Table 1**). In the intact sate, all cats performed 1:1 fore-hind patterns, with r values ranging from 0.73 to 0.98 (mean 0.93 ± 0.09) indicating consistent fore-hind coordination. At H1T1, we observed cycles with 1:1 patterns in all cats, with r values ranging from 0.33 to 0.96 (mean 0.68 ± 0.25) and six cats had cycles with 2:1 patterns, with r values ranging from 0.81 to 1.00 (mean 0.94 ± 0.06) and 0.64 to 0.99 (mean 0.87 ± 0.13) for the first and second forelimb cycles, respectively. At H1T2, we observed 1:1 patterns, with r values ranging from 0.33 to 0.97 (mean 0.75 ± 0.25) in six cats, while four cats had 2:1 patterns, with r values ranging from 0.76 to 0.99 (mean 0.90 ± 0.10) and 0.77 to 0.96 (mean 0.86 ± 0.08) for the first and second forelimb cycles, respectively. At H2T2, six cats had 1:1 patterns, with r values ranging from 0.15 to 0.84 (mean 0.58 ± 0.26) and 2:1 patterns, with r values ranging from 0.62 to 0.96 (mean 0.90 ± 0.05) and 0.68 to 0.81 (mean 0.71 ± 0.06) for the first and second forelimb cycles, respectively. Therefore, despite 2:1 patterns, which appear more consistent compared to 1:1 patterns after the first and second hemisections, coordination between the fore- and hindlimbs is maintained, albeit less consistent, after hemisections based on r values. It is important to note that cats received balance assistance after the second hemisection, which undoubtedly facilitated fore-hind coordination.

### Cutaneous reflexes after staggered hemisections in hindlimb muscles

To determine how cutaneous reflex pathways and transmission were affected by spinal lesions, we stimulated the left and right SP before and after the first (right T5-T6) and second (left T10-T11) hemisections in the same cats during quadrupedal treadmill locomotion and evaluated reflex responses in muscles of the four limbs and their potential role in locomotor recovery. Due to the large number of sampled muscles and because reflexes varied from one cat to another (Loeb, 1993; Frigon, 2011), out of 10 hindlimb muscles, we illustrate homonymous and crossed reflex responses in four muscles (SOL, VL, SRT and ST) bilaterally in representative cats. The SOL and VL muscles are mostly active during stance while SRT and ST are active during swing and/or at the stance-to-swing transition.

#### Homonymous responses

We stimulated the left and right SP nerves and recorded homonymous responses in muscles of the left and right hindlimbs, respectively, before and after staggered hemisections (**Fig. 5** and **Table 2**). **Figure 5** shows bilateral homonymous reflex responses in four phases in representative cats. For each state/time point, filled areas highlight evoked responses and are optimized for display according to the strongest response obtained in one of the four locomotor phases. The scale is optimized per state/time point and differs across state/time points. This is to show the pattern of evoked responses and its phase-dependent modulation at a given state/time point.

**Figure 5.**
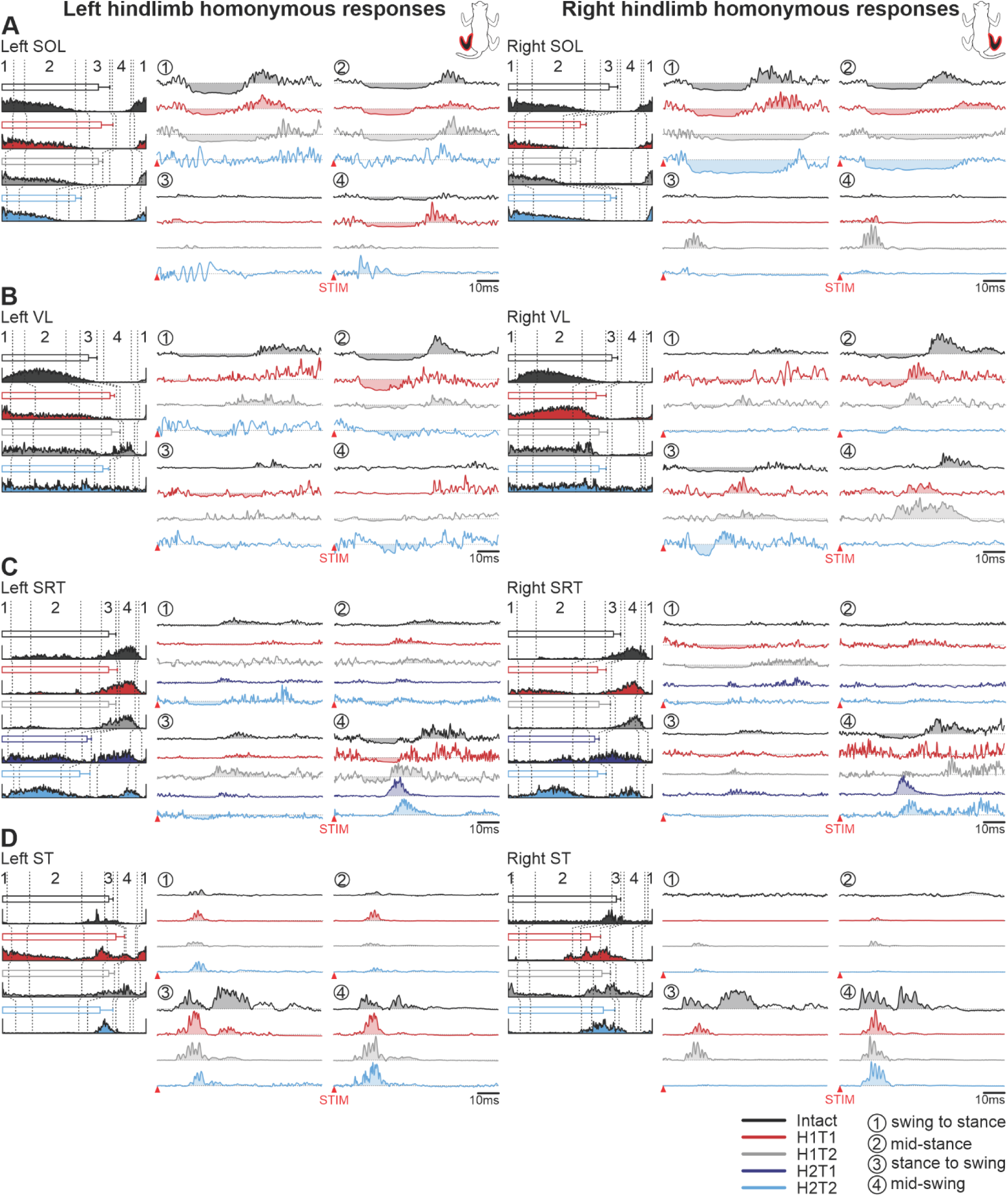
Phase-dependent modulation of cutaneous reflexes evoked in homonymous hindlimb muscles during locomotion before and following staggered hemisections. Each panel shows, from left to right, stance phases of the stimulated hindlimb (empty horizontal bars) with its averaged rectified muscle activity normalized to cycle duration in the different states/time points and homonymous reflex responses in representative cats for the left and right (**A**) soleus (SOL, cat AR), (**B**) vastus lateralis (VL, cat AR), (**C**) anterior sartorius (SRT, cat JA), and (**D**) semitendinosus (ST, cat GR). Reflex responses are shown with a post-stimulation window of 80 ms in four phases in the intact state, and after the first (H1) and second (H2) hemisections at time points 1 (T1) and/or 2 (T2). At each state/time point, evoked responses are scaled according to the largest response obtained in one of the four phases. The scale, however, differs between states/time points.

**Table 2.**
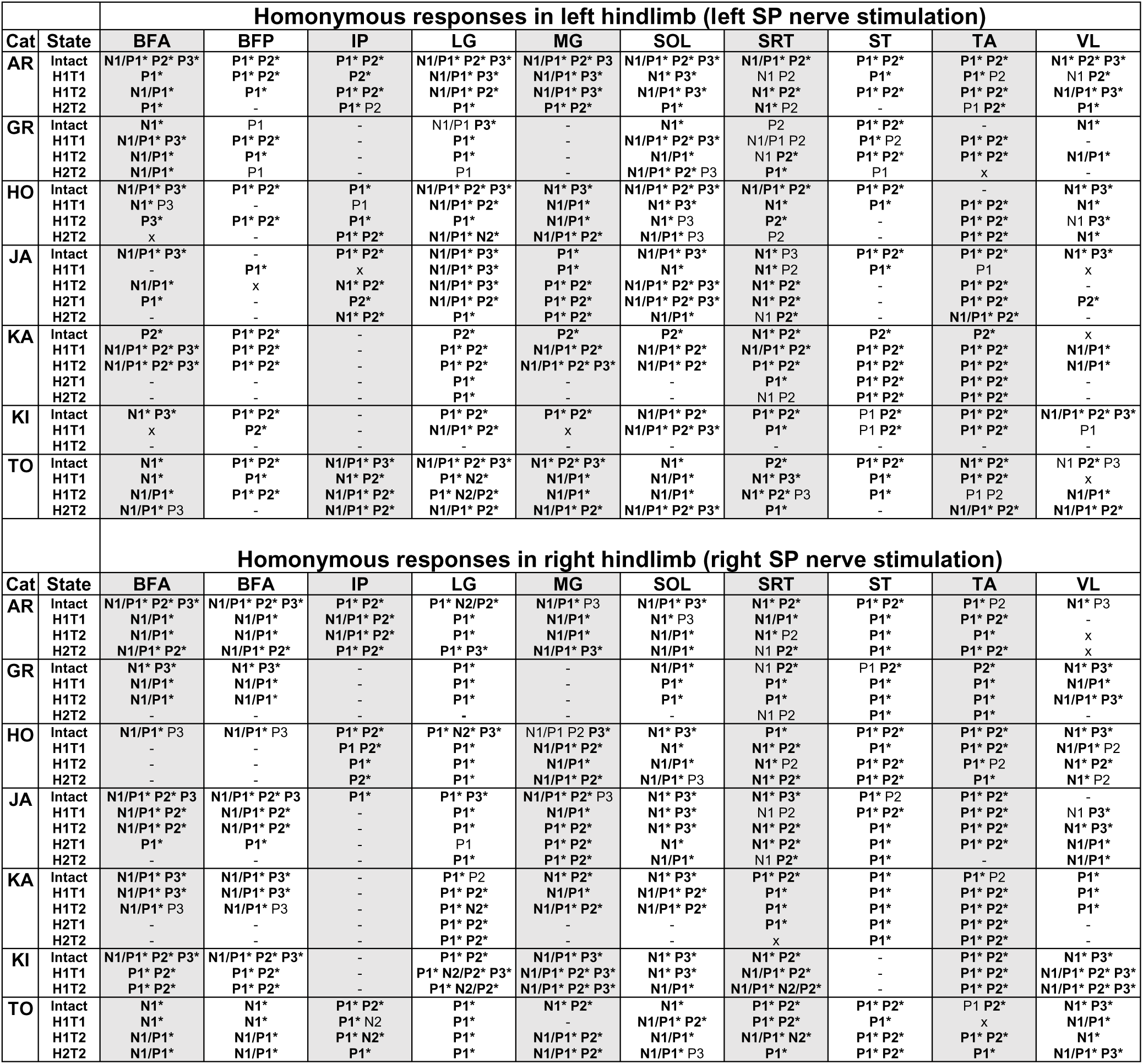
Homonymous reflex response occurrence before and after staggered hemisections. The table shows homonymous responses (P1, P2, P3, N1, N2 and N3) evoked in individual cats in left and right hindlimb muscles in the intact state, and after the first (H1) and second (H2) hemisections at 1-2 time points (T1 or T2). Asterisks indicate a significant phase modulation (one factor ANOVA, p < 0.05). x, No response. -, Non-implanted or non-analyzable (lost or excessive noise) muscle. BFA, biceps femoris anterior; BFP, biceps femoris posterior; IP, iliopsoas; LG, lateral gastrocnemius; MG, medial gastrocnemius; SOL, soleus; SRT, anterior sartorius; ST, semitendinosus; TA, tibialis anterior; VL, vastus lateralis.

In the left and right SOL (**Fig. 5A**), stimulating the SP nerve evoked homonymous N1 responses followed by P3 responses when the muscle was active (swing-to-stance and mid-stance) while weak P1 or N1 responses were observed at stance-to-swing and mid-swing in the intact state. After the first hemisection, N1 and P3 responses remained bilaterally at swing-to-stance and mid-stance at H1T1 but at H1T2 only N1 responses were observed in the right SOL. In the right SOL, homonymous P1 responses became prominent during mid-swing. After the second hemisection, at H2T2, reflex responses disappeared in the left SOL, despite a normal burst, while a prominent N1 was observed in the right SOL at swing-to-stance and mid-stance. We observed homonymous P1 responses at mid-swing bilaterally.

In the left and right VL (**Fig. 5B**), stimulating the SP nerve evoked homonymous N1 responses and/or P3 responses in all four phases in the intact state. After the first hemisection, at H1T1, we observed weaker and shorter duration N1 responses in some phases bilaterally while P3 responses disappeared bilaterally and P2 responses appeared, but in the right VL only. At H1T2, N1 responses were weak bilaterally, P2/P3 responses were present in the left VL at swing-to-stance and mid-stance and in all phases in the right VL. After the second hemisection, at H2T2, we observed N1 responses in left VL in all phases and no excitatory responses while in the right VL, we observed responses at the stance-to-swing transition only, with N1 followed by P2 responses.

In the left and right SRT (**Fig. 5C**), stimulating the SP nerve evoked homonymous N1 responses and/or P3 responses in all four phases in the intact state. The strongest responses, N1 followed by P3 responses, were found at-mid swing when the muscle had peak activity. After the first hemisection, at H1T1 and H1T2, responses in the right SRT disappeared at mid-swing while an N1 response remained in the left SRT and a P2 response appeared at H1T2. In the other phases, we observed weak N1 and P2 responses. After the second hemisection, at H2T1 and H2T2, we observed prominent P2 responses during mid-swing, with weak N1 and/or P2 responses in the other phases or no responses.

In the left and right ST (**Fig. 5D**), stimulating the SP nerve evoked homonymous P1 responses in most phases that could be followed by P2 responses, particularly at stance-to-swing and at mid-swing in the intact state. After the first and second hemisections, at H1T1, H1T2 and H2T2, P1 responses remained but P2 responses were lost or greatly reduced bilaterally. The P1 responses remained largest at mid-swing.

**Table 2** presents homonymous reflex response patterns observed in all 10 hindlimb muscles bilaterally for the 7 cats before and after staggered hemisections. Responses in bold with an asterisk indicate that they were significantly phase modulated. Some response patterns could change after the first or second hemisections, depending on the individual cat, but in general the phase-dependent modulation remained. Some notable changes include the loss of P2 and/or P3 responses in the right hindlimb, especially in LG, MG, BFA, BFP and ST. Thus, overall, short-latency excitatory and inhibitory responses remained after staggered hemisections while mid- and long-latency responses were reduced or lost in homonymous hindlimb muscles bilaterally.

#### Crossed responses

We stimulated the left and right SP nerves and recorded crossed responses in muscles of the right and left hindlimbs, respectively, before and after staggered hemisections in the four phases (**Fig. 6** and **Table 3**). Although the four phases are defined according to the stimulated limb, it is important to consider the phase of the contralateral limb where the muscles are recorded.

**Figure 6.**
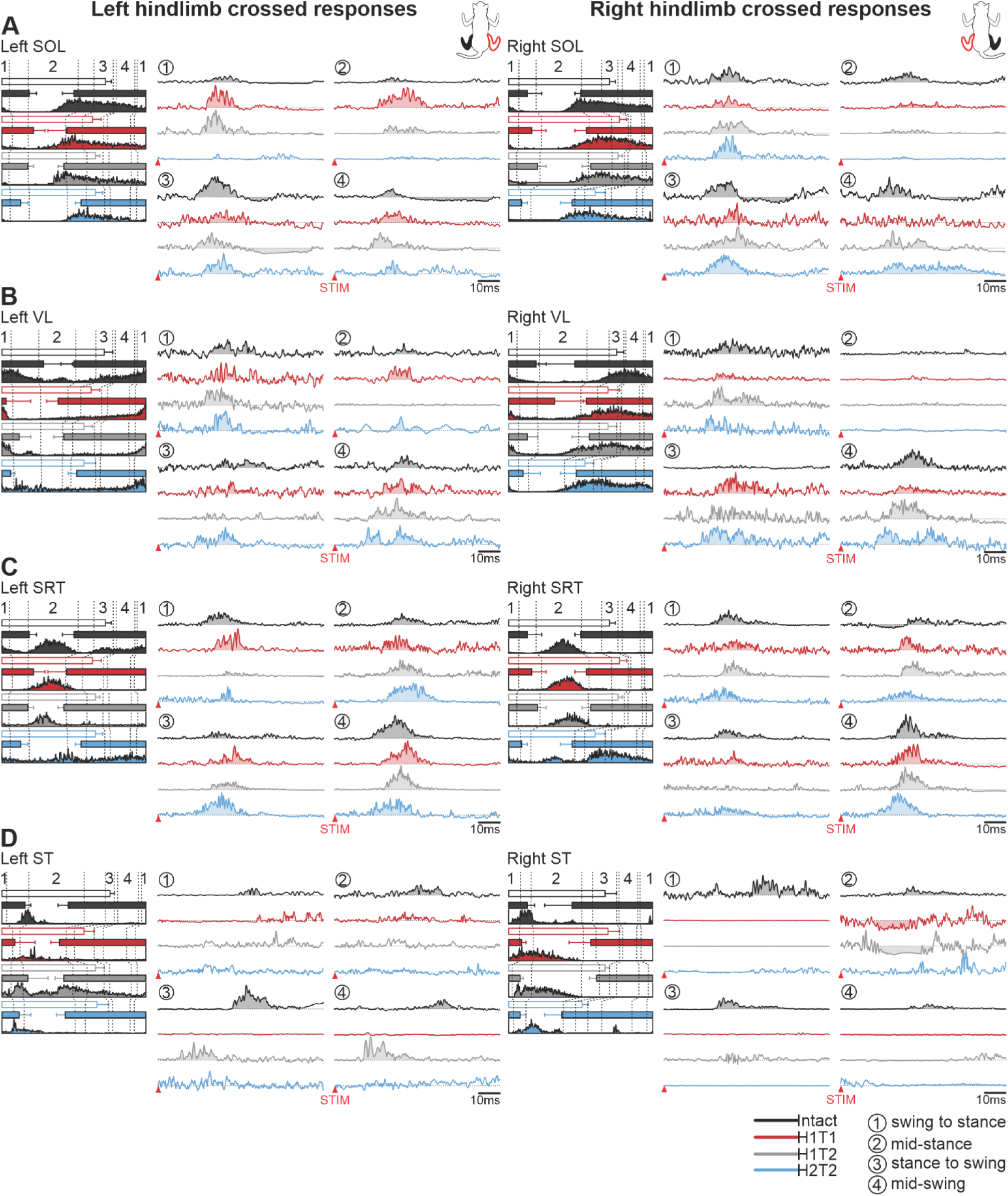
Phase-dependent modulation of cutaneous reflexes evoked in crossed hindlimb muscles during locomotion before and following staggered hemisections. Each panel shows, from left to right, stance phases of the stimulated hindlimb (empty horizontal bars) and crossed hindlimb (filled horizontal bars) with its averaged rectified muscle activity normalized to cycle duration in the different states/time points and crossed reflex responses in representative cats for the left and right (**A**) soleus (SOL, cat HO), (**B**) vastus lateralis (VL, cat TO), (**C**) anterior sartorius (SRT, cat HO), and (**D**) semitendinosus (LST, cat GR; RST, cat AR). Reflex responses are shown with a post-stimulation window of 80 ms in four phases in the intact state, and after the first (H1) and second (H2) hemisections at time points 1 (T1) and/or 2 (T2). At each state/time point, evoked responses are scaled according to the largest response obtained in one of the four phases. The scale, however, differs between states/time points.

**Table 3.**
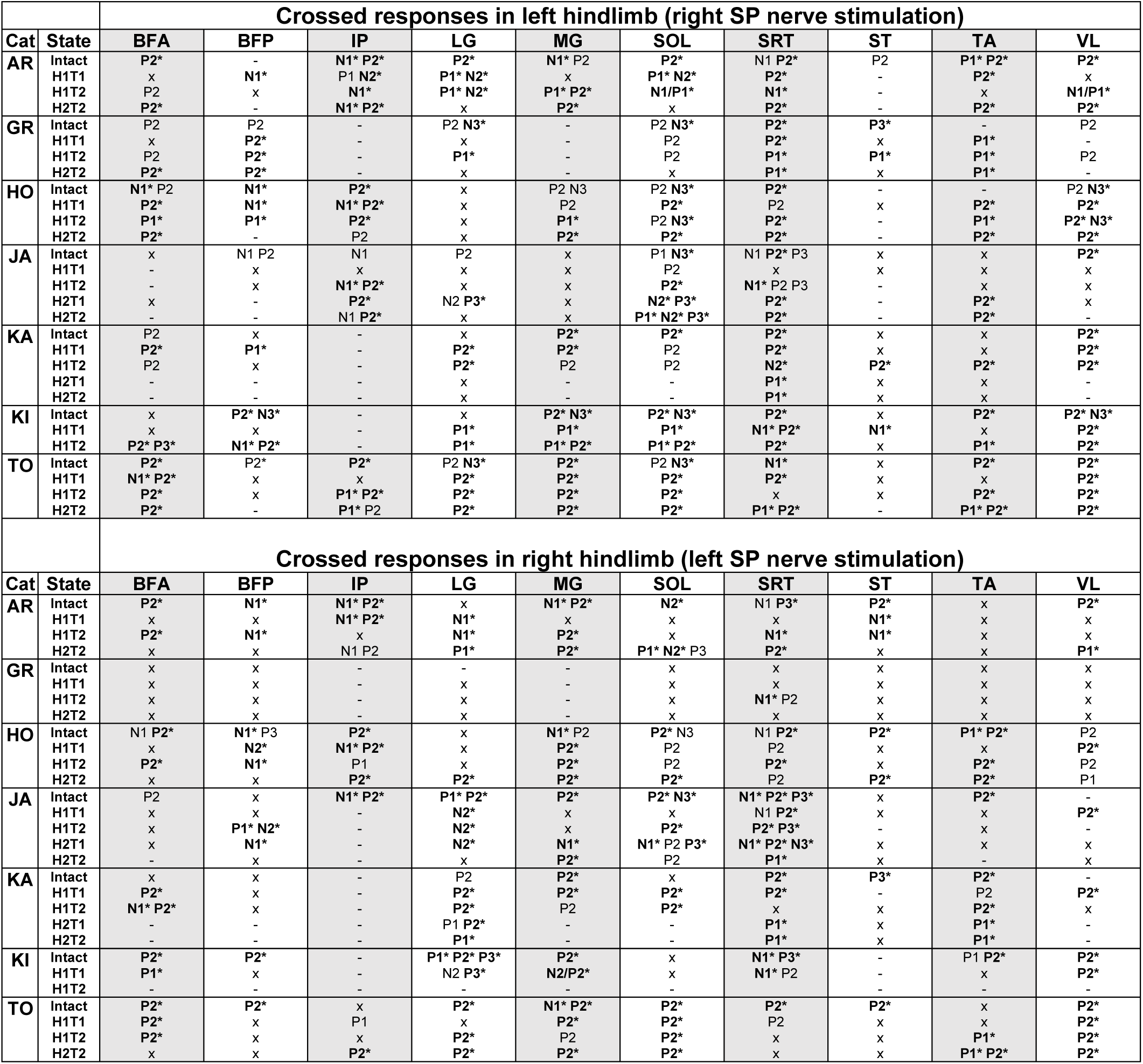
Crossed reflex responses before and after staggered hemisections. The table shows crossed responses (P1, P2, P3, N1, N2 and N3) evoked in individual cats in left and right hindlimb muscles in the intact state, and after the first (H1) and second (H2) hemisections at 1-2 time points (T1 or T2). Asterisks indicate a significant phase modulation (one factor ANOVA, p < 0.05). x, No response. -, Non-implanted or non-analyzable (lost or excessive noise) muscle. BFA, biceps femoris anterior; BFP, biceps femoris posterior; IP, iliopsoas; LG, lateral gastrocnemius; MG, medial gastrocnemius; SOL, soleus; SRT, anterior sartorius; ST, semitendinosus; TA, tibialis anterior; VL, vastus lateralis.

In the left and right SOL (**Fig. 6A**), stimulating the SP nerve in the intact state evoked crossed P2 responses followed by N3 responses in all four phases that were most prominent when the muscle was active, at stance-to-swing and mid-swing of the stimulated limb. After the first hemisection, at H1T1 and H1T2, P2 responses remained in all four phases, with the largest responses observed when the muscle was active (stance-to-swing and mid-swing of the stimulated limb). We observed that N3 responses disappeared at H1T1 before reappearing at H1T2, but only in the left SOL during stance to swing of the stimulated limb. After the second hemisection, at H2T2, P2 responses were present bilaterally in all phases, except at mid-stance of the stimulated limb when the muscle was inactive.

In the left and right VL (**Fig. 6B**), stimulating the SP nerve in the intact state evoked crossed P2 responses particularly during the muscles’ activity (mid-swing and swing-to-stance of the stimulated limb). After the first and second hemisections, at H1T1, H1T2 and H2T2, P2 responses remained prominent bilaterally when VL was active and during its inactivity (mid-stance of the stimulated limb) for the left VL, except at H1T2.

In the left and right SRT (**Fig. 6C**), stimulating the SP nerve in the intact state evoked crossed P2 responses in all four phases that were most prominent when the muscle was inactive (mid-swing of the stimulated limb). In the right VL, P2 responses were also preceded by N1 responses during its activity (mid-stance of the stimulated limb). After the first and second hemisections, at H1T1, H1T2 and H2T2, we observed large P2 responses bilaterally during mid-swing of the stimulated limb, with weaker responses in most other phases.

In the left and right ST (**Fig. 6D**), stimulating the SP nerve in the intact state evoked crossed P2/P3 responses in all four phases. After the first hemisection, at H1T1, P2 responses disappeared bilaterally while strong N1 responses appeared in the right ST during mid-stance of the stimulated limb. At H1T2, P1 responses appeared in the left ST during stance-to-swing and mid-swing of the stimulated limb, while N1 responses remained in the right ST. After the second hemisection, at H2T2, no crossed responses were evoked bilaterally.

**Table 3** shows crossed reflex response patterns observed in all 10 hindlimb muscles bilaterally for the 7 cats before and after staggered hemisections. Although some response patterns changed after the first or second hemisections, the phase-dependent modulation generally remained. Some notable changes in crossed reflex response patterns include a greater number of P2 responses disappearing in the right hindlimb (BFP, BFA) compared to the left, and a loss of long-latency P3/N3 responses bilaterally (SOL and SRT).

### Cutaneous reflexes before and after staggered hemisections in forelimb muscles

We recorded from 5 forelimb muscles and we illustrate homolateral and diagonal reflex responses evoked by stimulating the SP nerves in 3 muscles (ECU, TRI and BB) bilaterally in representative cats. The ECU and TRI muscles are mostly active during stance while BB is active during swing and/or at the stance-to-swing transition. As with homonymous and crossed responses in hindlimb muscles, the four phases are defined according to the stimulated limb. It is thus important to consider the phase of the forelimb where the muscles are recorded.

#### Homolateral responses

We stimulated the left and right SP nerves and recorded homolateral responses in muscles of the left and right forelimbs, respectively, before and after staggered hemisections (**Fig. 7** and **Table 4**).

**Figure 7.**
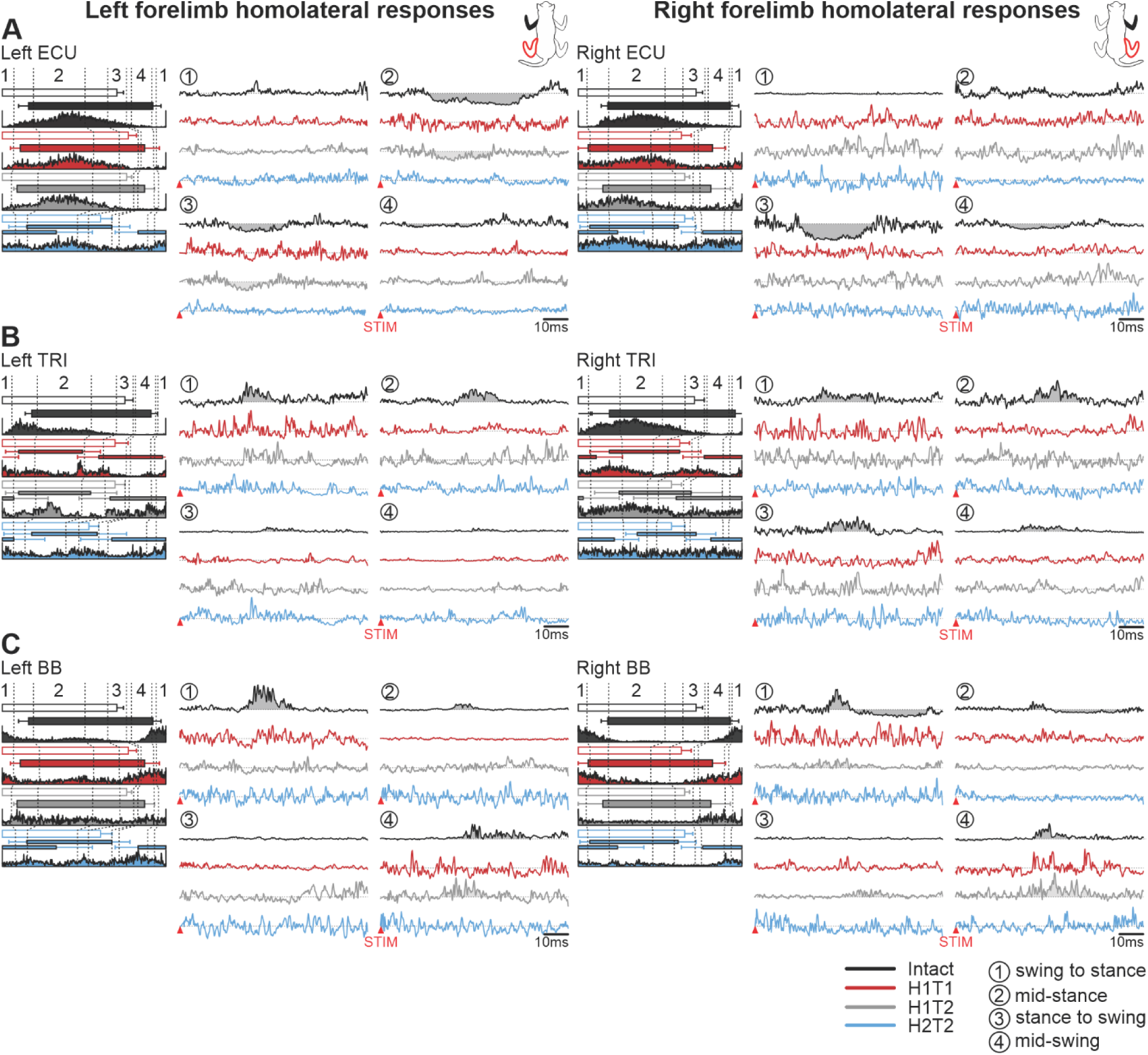
Phase-dependent modulation of cutaneous reflexes evoked in homolateral forelimb muscles during locomotion before and following staggered hemisections. Each panel shows, from left to right, stance phases of the stimulated hindlimb (empty horizontal bars) and homolateral forelimb (filled horizontal bars) with its averaged rectified muscle activity normalized to cycle duration in the different states/time points and homolateral reflex responses in representative cats for the left and right (**A**) extensor carpi ulnaris (ECU, cat HO), (**B**) triceps brachii (TRI, cat TO), and (**C**) biceps brachii (BB, cat HO). Reflex responses are shown with a post-stimulation window of 80 ms in four phases in the intact state, and after the first (H1) and second (H2) hemisections at time points 1 (T1) and/or 2 (T2). At each state/time point, evoked responses are scaled according to the largest response obtained in one of the four phases. The scale, however, differs between states/time points.

**Table 4.**
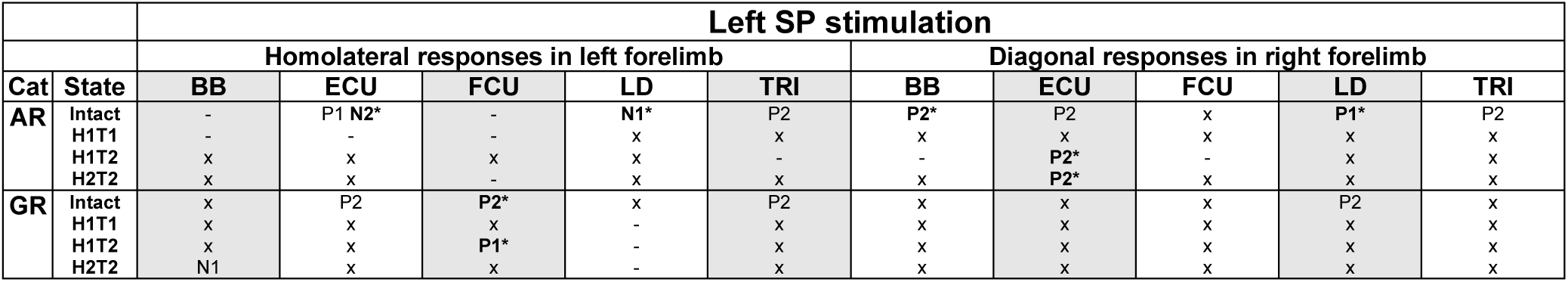
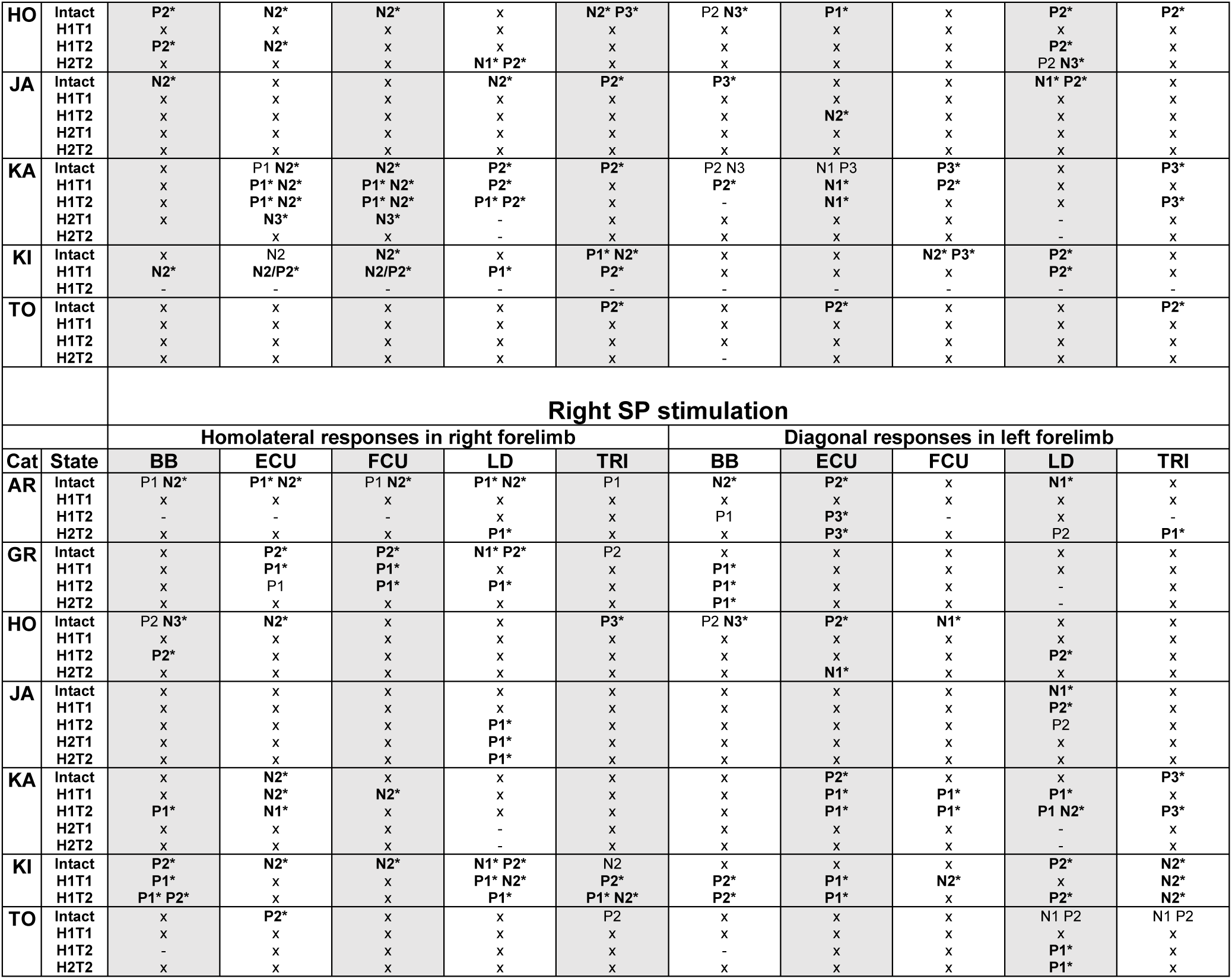
Homolateral and diagonal reflex responses before and after staggered hemisections. The table shows homolateral and diagonal responses (P1, P2, P3, N1, N2 and N3) evoked in individual cats in left and right forelimb muscles in the intact state, and after the first (H1) and second (H2) hemisections at 1-2 time points (T1 or T2). Asterisks indicate a significant phase modulation (one factor ANOVA, p < 0.05). x, No response. -, Non-implanted or non-analyzable (lost or excessive noise) muscle. BB, biceps brachii; ECU, extensor carpi ulnaris; FCU, flexor carpi ulnaris; LD, latissimus dorsi; TRI, triceps brachii.

In the left and right ECU (**Fig. 7A**), stimulating the SP nerve in the intact state evoked homolateral N2 responses mostly when the muscle was active or at the end of its activity (mid-stance and stance-to-swing of the stimulated hindlimb). After the first hemisection, at H1T1, no responses were observed bilaterally while N2 responses reappeared on the left side at H1T2. After the second hemisection, at H2T2, stimulation of the left or right SP did not evoke responses.

In the left and right TRI (**Fig. 7B**), stimulating the SP nerve in the intact state evoked homolateral P2 responses in all four phases that were most prominent at the muscle’s peak activity (mid-stance of the stimulated hindlimb). After the first and second hemisections, at H1T1, H1T2 and H2T2, P2 responses completely disappeared bilaterally, and we observed no other responses.

In the left and right BB (**Fig. 7C**), stimulating the SP nerve in the intact state evoked homolateral P2 responses bilaterally followed by N3 responses but only for the right side in this cat. The most prominent P2 and N3 responses were observed when the muscle was active (swing-to-stance of stimulated hindlimb).

After the first hemisection, at H1T1, no responses were observed bilaterally but P2 responses reappeared at H1T2 when the muscle was active (mid-swing of stimulated hindlimb). After the second hemisection, at H2T2, stimulation of the left or right SP did not evoke responses.

#### Diagonal responses

We stimulated the left and right SP nerves and recorded diagonal responses in muscles of the right and left forelimbs, respectively, before and after staggered hemisections (**Fig. 7** and **Table 4**).

In the left and right ECU (**Fig. 8A**), stimulating the SP nerve in the intact state evoked diagonal P2 responses in all four phases that were most prominent when the muscle was active (mid-swing and swing-to-stance of the stimulated hindlimb). After the first hemisection, at H1T1, P2 responses disappeared bilaterally. At H1T2, P2 or P3 responses appeared bilaterally and were generally present in all four phases. The strongest responses were found at swing-to-stance or mid-swing of the stimulated hindlimb in the left and right ECU, respectively. After the second hemisection, at H2T2, P2 and P3 diagonal responses remained in the right and left ECU, respectively.

**Figure 8.**
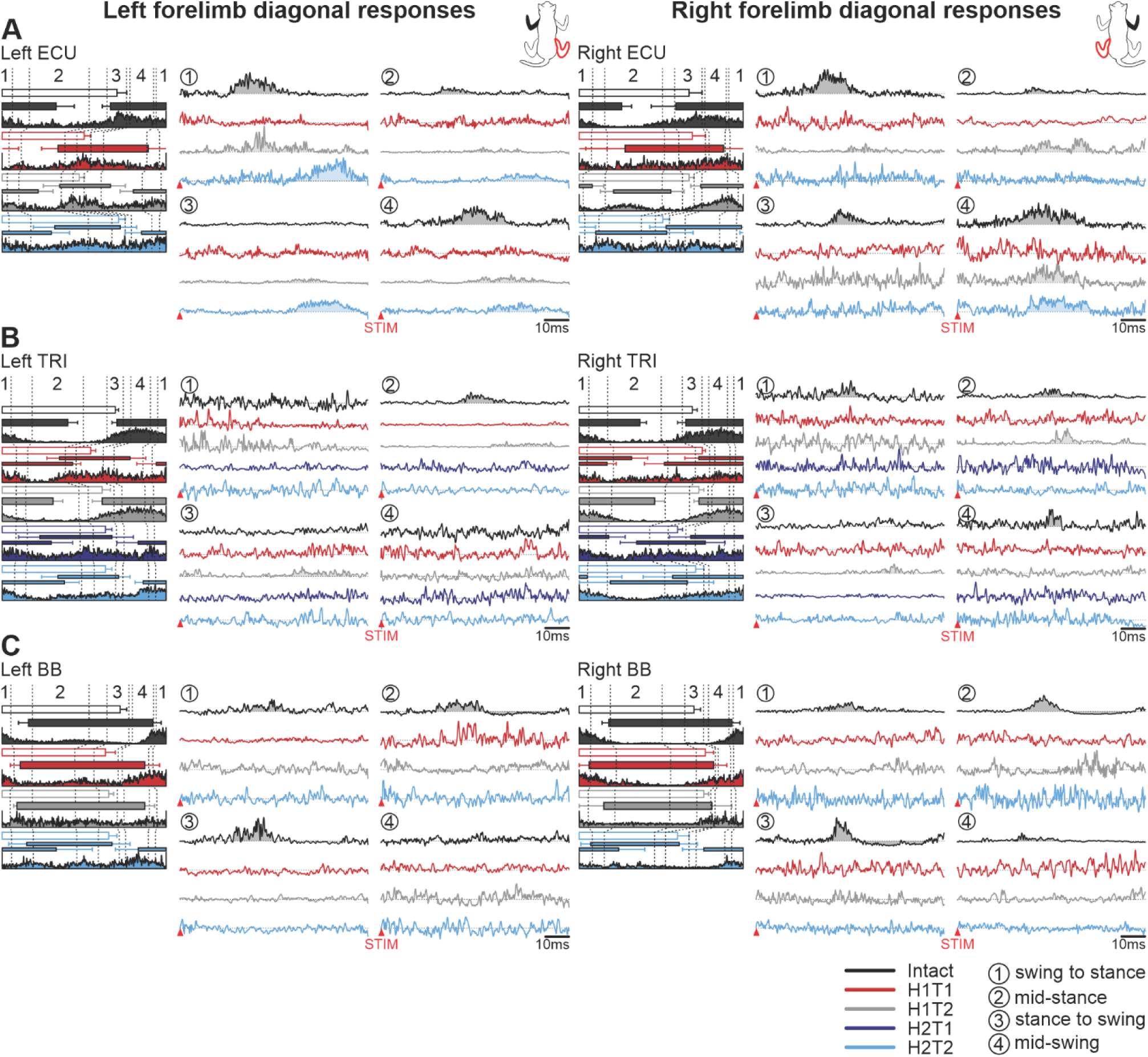
Phase-dependent modulation of cutaneous reflexes evoked in diagonal forelimb muscles during locomotion before and following staggered hemisections. Each panel shows, from left to right, stance phases of the stimulated hindlimb (empty horizontal bars) and diagonal forelimb (filled horizontal bars) with its averaged rectified muscle activity normalized to cycle duration in the different states/time points and diagonal reflex responses in representative cats for the left and right (**A**) extensor carpi ulnaris (ECU, cat AR), (**B**) triceps brachii (TRI, cat KA), and (**C**) biceps brachii (BB, cat HO). Reflex responses are shown with a post-stimulation window of 80 ms in four phases in the intact state, and after the first (H1) and second (H2) hemisections at time points 1 (T1) and/or 2 (T2). At each state/time point, evoked responses are scaled according to the largest response obtained in one of the four phases. The scale, however, differs between states/time points.

In the left and right TRI (**Fig. 8B**), stimulating the SP nerve in the intact state evoked diagonal P3 responses mostly when the muscle was active (mid-stance of the stimulated hindlimb). After the first hemisection, at H1T1, P3 responses disappeared bilaterally then reappeared at H1T2 at mid-stance and stance-to swing of the stimulated hindlimb. After the second hemisection, at H2T2 and H2T2, no diagonal responses were evoked bilaterally.

In the left and right BB (**Fig. 8C**), stimulating the SP nerve in the intact state evoked diagonal P2 responses bilaterally followed by N3 responses. We observed the most prominent P2 and N3 responses during the muscles’ inactivity, when the stimulated hindlimb was in mid-stance and at the stance-to-swing transition. After the first and second hemisections, at H1T1, H1T2 and H2T2, P2 and N3 diagonal responses disappeared bilaterally and no other responses were evoked.

**Table 4** shows homolateral and diagonal reflex response patterns observed in all 5 forelimb muscles bilaterally for the 7 cats before and after staggered hemisections. Although the phase-dependent modulation of responses generally remained after staggered hemisections, when responses were present, we observed a loss in response occurrence in most muscles after the first and/or second hemisections.

### Staggered hemisections reduce the occurrence of evoking mid- and long-latency responses

After complete or incomplete spinal lesions in cats, mid- and long-latency responses in hindlimb muscles are generally reduced or abolished (Fuwa *et al*., 1991; LaBella *et al*., 1992; Frigon & Rossignol, 2008; Frigon *et al*., 2009; Hurteau *et al*., 2017). Here, we investigated the probability of evoking reflex responses in all four limbs before and after staggered hemisections by evaluating the distribution of SLRs (N1/P1), MLRs (N2/P2) and LLRs (N3/P3). We did this by calculating the fraction of the total number of SLRs or MLRs/LLRs separately, on recorded muscles for each limb across cats. We excluded the H2T1 time point as only two cats were recorded.

We found no significant difference in response occurrence probability for left and right homonymous SLRs across states/time points (**Fig. 9A**). However, we found a significant main effect of state/time point for left (p = .008, GLMM) and right (p = 4.50 × 10^-5^, GLMM) homonymous MLRs/LLRs. Left homonymous MLRs/LLRs in the intact state were 4.6 (p = .001), 3.1 (p = .019) and 4.0 (p = .006) times more likely to be evoked compared to H1T1, H1T2 and H2T2, respectively. For right homonymous MLRs/LLRs, they were 6.9 (p = 2.50 × 10^-5^), 7.4 (p = 1.20 × 10^-5^) and 4.8 (p = .001) times more likely to be evoked compared to H1T1, H1T2 and H2T2, respectively. The probability of evoking crossed SLRs in left or right hindlimb muscles as well as crossed MLR/LLRs in the left hindlimb did not differ across states/time points (**Fig. 9B**). However, the probability of evoking crossed MLRs/LLRs in right hindlimb muscles was significantly affected by state/time point (p = .003, GLMM), with 4.1 (p = .001), 3.8 (p = .004) and 4.2 (p = .002) more chance of being evoked in the intact state compared to H1T1, H1T2 and H2T2, respectively.

**Figure 9.**
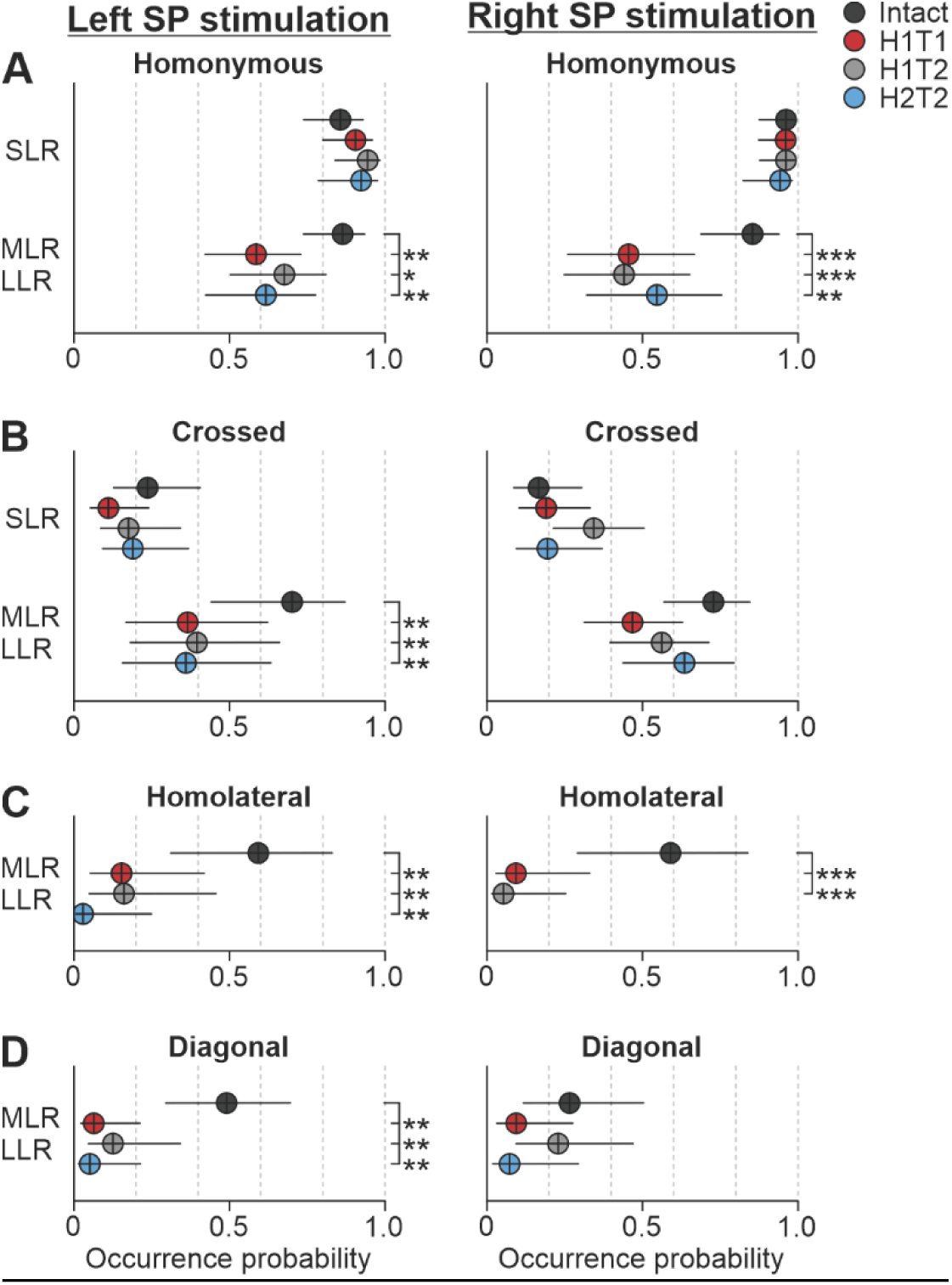
Reflex response occurrence in all four limbs before and after staggered hemisections. Response occurrence probabilities are shown for short- (SLR) and mid-/long-latency (MLRs/LLRs) responses with stimulation of the left or right superficial peroneal nerve before (intact) and after the first (H1) and second (H2) hemisections at time points 1 (T1) and/or 2 (T2). Tables 2 and 3 provide details on the number of pooled data for SLRs, MLRs and LLRs. Each filled circle represents the mean probability ± confidence interval (95%) in 10 hindlimb or 5 forelimb muscles pooled across cats for homonymous/crossed (**A** and **B**) and homolateral/diagonal (**C** and **D**) responses, respectively. If a significant main effect of state/timepoint was found (generalized linear mixed model), we compared states/time points. Asterisks indicate significant differences at p < 0.05*, p < 0.01** and p < 0.001***. When one state/time point was significantly different from two states/time points, the comparison starts with a longer horizontal line.

For homolateral (**Fig. 9C**) and diagonal (**Fig. 9D**) responses in the left and right forelimbs, we only evaluated MLRs/LLRs because of weaker occurrence and fewer sampled muscles. We found a significant main effect of state/time point on homolateral (p = 4.30 × 10^-4^, GLMM) and diagonal (p = 2.29 × 10^-4^, GLMM) MLRs/LLRs occurrence probability with left SP nerve stimulation and in homolateral (p = 1.87 × 10^-4^, GLMM) MLRs/LLRs with right SP nerve stimulation. Responses were always more likely to be evoked in the intact state compared to after the first and second hemisections. Note that no diagonal responses were evoked in the left forelimb with right SP nerve stimulation. Homolateral responses in the left forelimb were 7.8 (p = .002), 7.6 (p = .004) and 45.5 (p = .001) times more likely to be evoked in the intact state compared to H1T1, H1T2 and H2T2, respectively. Diagonal responses in the right forelimb were 13.7 (p = .001), 6.5 (p = .008) and 17.5 (p = .001) times more likely to be evoked in the intact state compared to H1T1, H1T2 and H2T2, respectively. Homolateral responses in the right forelimb were 13.5 (p = 2.60 × 10^-4^) and 25.0 (p = 1.70 × 10^-4^) times more likely to be evoked in the intact state compared to H1T1 and H1T2, respectively. We observed no difference between states/time points in diagonal MLR/LLR of the left forelimb likely because occurrence probability was low in the intact state.

Therefore, overall, after the first or second hemisection, the probability of evoking MLRs/LLRs in all four limbs with both left and right SP nerve stimulation is generally always lower in all four limbs compared to the intact state. The only exceptions were for crossed and diagonal MLRs/LLRs in the left hindlimb (p = .051, GLMM) and left forelimb (p = .136, GLMM), respectively.

### Temporal characteristics of hindlimb cutaneous reflexes change after staggered hemisections

To determine whether temporal characteristics of homonymous responses were affected after the first and second hemisections, we measured latencies for SLRs and MLRs/LLRs, as well as durations of short-latency inhibitory responses. We excluded the H2T1 state/time point as only two cats were recorded.

#### Latencies

The latency of reflex responses provides an estimate of the number of synaptic relays in the pathway. We measured the latencies of homonymous SLRs by pooling N1 and P1 responses for the left and right hindlimbs (**Fig. 10A**). We found no significant differences in left SLR latencies in the left hindlimb (p = .128, LMM). In the right hindlimb, we found a main effect of state/time point (p = .043, LMM) with significantly longer SLRs at H1T1 compared to the intact state (p = .046) and H1T2 (p = .012). For homonymous MLRs/LLRs in the left (p = .060, LMM) and right (p = .213, LMM) hindlimbs (**Fig. 10B**), we found no effect of state/time point on their latencies.

**Figure 10.**
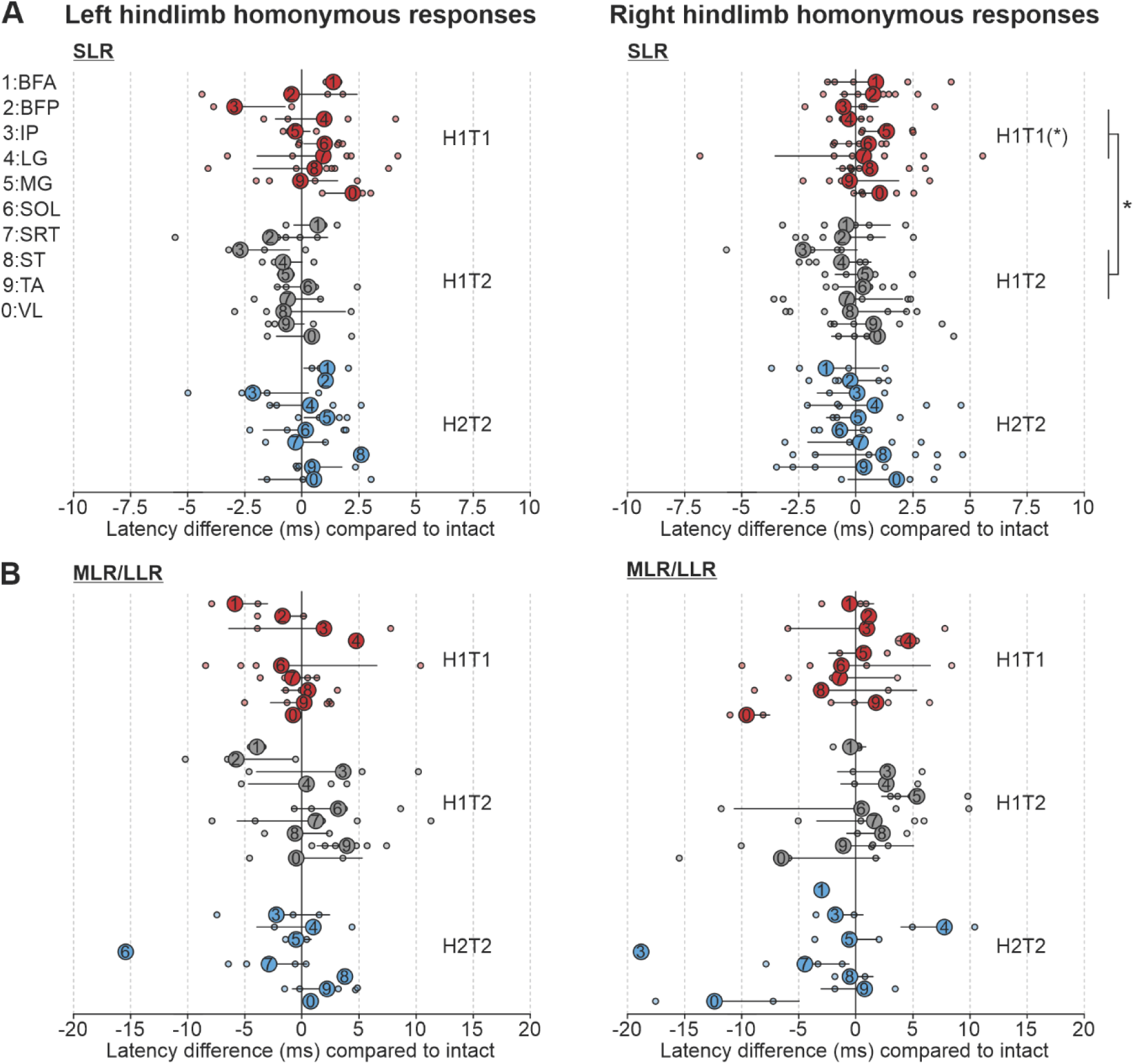
Latencies of homonymous reflex responses before and after staggered hemisections. Each panel shows changes in homonymous response latencies in different muscles compared to the intact state for (**A**) short-latency (SLRs) and (**B**) mid-/long-latency (MLRs/LLRs) responses in left and right hindlimb muscles after the first (H1) and second (H2) hemisections at time points 1 (T1) and/or 2 (T2). Large circles represent the mean ± SD while small circles show individual data points. When a significant main effect of state/time point was found (linear mixed model), we compared states/time points. Asterisks in parentheses indicate a significant difference from the intact state while other asterisks indicate significant differences between states/time points at *p < 0.05. BFA, biceps femoris anterior; BFP, biceps femoris posterior; IP, iliopsoas; LG, lateral gastrocnemius; MG, medial gastrocnemius; SOL, soleus; SRT, anterior sartorius; ST, semitendinosus; TA, tibialis anterior; VL, vastus lateralis.

#### Durations

We measured the duration of homonymous N1 responses by pooling responses in extensors (BFA, LG, MG, SOL, and VL) of the left and right hindlimbs (**Table 5**). To compare states/time points, we pooled durations of the different hindlimb extensors obtained across cats. We found a significant main effect of state/time point (p = .001, LMM) in left hindlimb extensors. N1 durations were significantly reduced at H1T1 (p = .022) and H2T2 (p = 2.23 × 10^-4^) compared to the intact state, and at H1T1 (p = .033) and H2T2 (p = 3.93 × 10^-4^) compared to H1T2. We observed no significant difference for right hindlimb extensors (p = .084, LMM).

**Table 5.** Short-latency inhibitory response durations in homonymous extensors before and after staggered hemisections. The table shows the mean (± SD) durations of N1 responses in left (L) and right (R) hindlimb extensors, for the first hemisection (H1) at two time points (T1/T2) and at the second time point (T2) after the second hemisection (H2). The number in brackets indicates the number of cats. BFA, biceps femoris anterior; LG, lateral gastrocnemius; MG, medial gastrocnemius; SOL, soleus; VL, vastus lateralis.

|  |  | Intact | H1T1 | H1T2 | H2T2 |
| --- | --- | --- | --- | --- | --- |
| Left SP stimulation | LBFA | 33.5 ± 3.9 ms (6) | 27.4 ± 7.5 ms (4) | 32.0 ± 6.1 ms (5) | 27.7 ± 6.1 ms (2) |
|  | LLG | 26.3 ± 3.1 ms (4) | 23.8 ± 8.4 ms (4) | 24.6 ± 4.5 ms (3) | 23.6 ± 1.9 ms (2) |
|  | LMG | 27.0 ± 9.7 ms (3) | 25.4 ± 3.6 ms (3) | 29.1 ± 6.7 ms (3) | 18.9 ± 1.2 ms (2) |
|  | LSOL | 39.5 ± 3.8 ms (6) | 31.1 ± 7.7 ms (7) | 38.3 ± 6.4 ms (6) | 26.5 ± 4.1 ms (4) |
|  | LVL | 28.2 ± 6.4 ms (6) | 28.2 ± 4.7 ms (4) | 29.6 ± 8.4 ms (5) | 20.3 ± 12.1 ms (2) |
|  | **Total mean** | **30.9 ms** | **27.2 ms** | **30.7 ms** | **23.4 ms** |
| Right SP stimulation | RBFA | 30.9 ± 5.3 ms (7) | 33.5 ± 7.7 ms (5) | 33.5 ± 6.4 ms (5) | 31.8 ± 10.8 ms (2) |
|  | RMG | 27.3 ± 9.7 ms (4) | 26.9 ± 9.9 ms (3) | 35.3 ± 16.5 ms (3) | 22.6 ± 9.6 ms (3) |
|  | RSOL | 36.9 ± 9.5 ms (7) | 36.5 ± 8.5 ms (6) | 40.3 ± 10.3 ms (6) | 37.9 ± 8.1 ms (4) |
|  | RVL | 31.9 ± 3.2 ms (5) | 25.2 ± 9.2 ms (5) | 28.1 ± 7.5 ms (5) | 23.0 ± 5.9 ms (3) |
|  | **Total mean** | **31.8 ms** | **30.5 ms** | **34.3 ms** | **28.8 ms** |

## DISCUSSION

In the present study, we showed changes in reflex responses evoked by electrically stimulating cutaneous afferents of the foot dorsum (SP nerve stimulation) during locomotion after staggered hemisections. Changes in reflex responses included a loss/reduction of mid and long-latency responses in all four limbs after the first and second hemisections, particularly when the stimulation is delivered to the left SP. The durations of short-latency inhibitory responses in left ipsilateral extensors were also significantly shorter early after the first hemisection, before returning towards intact values and reduced again after the second hemisection. These many changes in reflex responses correlated with altered fore-hind coordination and impaired balance during quadrupedal locomotion. The main results are summarized in **Table 6**. We discuss how these findings provide insight on neural pathways and locomotor control/recovery after SCI.

**Table 6.**
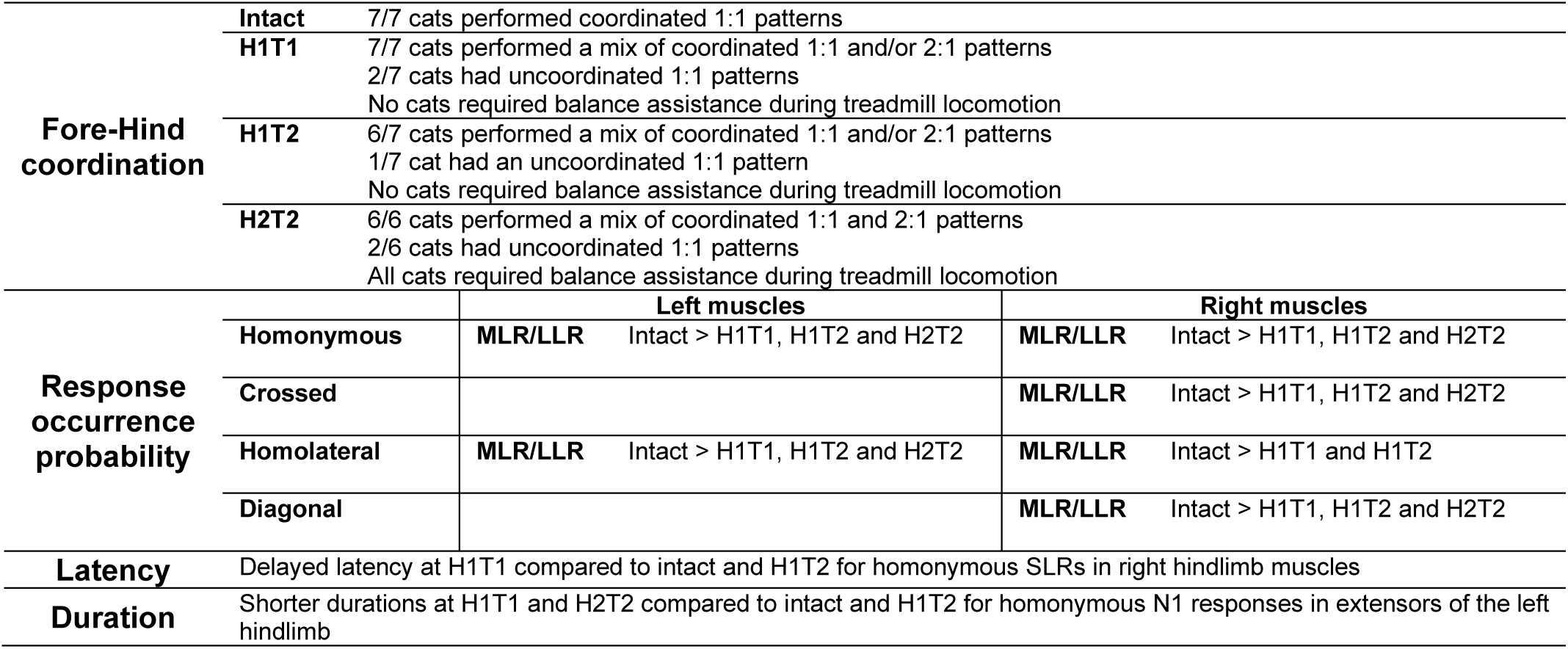
Summary of main results.

### Altered coordination of the fore- and hindlimbs and impaired dynamic balance after staggered hemisections

All cats, except one, performed cycles with 2:1 fore-hind patterns after spontaneously recovering quadrupedal locomotion after the first and second hemisections (**Table 1**), as shown previously after a single lateral hemisection (Barrière *et al*., 2010; Thibaudier *et al*., 2017) or with staggered hemisections (Kato *et al*., 1984, 1985; Audet *et al*., 2023). Although fore-hind coordination remained relatively consistent on a step-by-step basis based on circular statistics, three cats (HO, JA and TO) showed impaired coordination (r values below 0.50) at one or more time points after hemisections, mainly with 1:1 patterns, which did not appear to correlate with lesion extent (**Fig. 3**). Studies have reported that lesion extent rather than the loss of a particular spinal pathway is responsible for 2:1 patterns (English, 1980, 1985; Kloos *et al*., 2005; Górska *et al*., 2013). Another possible explanation for the 2:1 patterns may result from reduced inhibition from the hindlimb central pattern generators (CPGs) to the forelimb CPGs (Budakova & Shik, 1974; Górska *et al*., 2013; Thibaudier *et al*., 2017), thereby increasing forelimb CPG excitability and/or rhythmicity. Surprisingly, circular statistics showed that r values were generally higher with 2:1 patterns compared to 1:1 patterns after hemisections, consistent with greater step-by-step consistency. This suggests that adopting 2:1 patterns is a compensatory strategy to improve fore-hind coordination and possibly stability/balance. Intact cats also perform 2:1 patterns on a transverse split-belt treadmill when the forelimbs step faster than the hindlimbs (Frigon *et al*., 2014; Thibaudier & Frigon, 2014; Thibaudier *et al*., 2017). Functionally, performing more and shorter forelimb steps could maximize stability by shifting the center of gravity rostrally and having it more centered within the support polygon (Cartmill *et al*., 2002), as well as preventing interference between homolateral limbs (Lecomte *et al*., 2022; Audet *et al*., 2023).

An important caveat is that balance assistance was required after the second hemisection to conduct reflex testing during locomotion, with an experimenter holding the tail for stability, but without providing weight support. Without this balance assistance, cats stumbled and fell every few steps. Thus, consistent fore-hind coordination after the second hemisection was only possible with balance assistance. Other studies that reported impaired fore-hind coordination did not provide balance assistance (Kato *et al*., 1984; Murray *et al*., 2010; Cowley *et al*., 2015). Indeed, a main challenge during walking after SCI in cats and humans is controlling posture/balance (Fouad & Pearson, 2004; Van Hedel & Dietz, 2010; Rossignol & Frigon, 2011). Spinal-transected cats can support their weight during quiet standing but cannot maintain balance when moving or when perturbed (Pratt *et al*., 1994; Fung & Macpherson, 1999; Macpherson & Fung, 1999). Supraspinal structures are required for dynamic balance (reviewed in Deliagina et al., 2014).

### Changes in reflex responses after staggered lateral hemisections and neural pathways involved

Staggered lateral hemisections disrupted descending and ascending spinal pathways first unilaterally (right side) and then bilaterally, with the magnitude of this disruption depending on lesion extent (**Fig. 3**). These lesions and the occurrence of reflex responses (**Tables 2-4** and **Figure 9**) allow us to predict putative neural pathways/mechanisms involved in short-, mid-, and long-latency responses. For instance, short-latency N1 and P1 responses in muscles of the ipsilateral (homonymous) and contralateral (crossed) hindlimb remained after the first and second hemisections with both left and right SP nerve stimulations. They also maintained their phase-dependent modulation. This indicates that these pathways are mainly mediated and modulated within lumbosacral circuits, although we cannot exclude some supraspinal/cervical contributions. Studies have shown that phase-dependent short-latency reflex responses remain following thoracic spinal transection, although some response patterns are altered (Forssberg *et al*., 1977; LaBella *et al*., 1992; Frigon & Rossignol, 2008; Hurteau *et al*., 2017; Hurteau & Frigon, 2018). Although N1 responses in homonymous extensor muscles still occurred after the first and second hemisections, their duration was considerably shortened early after the first hemisection (i.e., at H1T1) in the left hindlimb before returning towards intact values later on (i.e. at H1T2), and then shortened once again after the second hemisection (**Table 5**). Thus, the loss of descending pathways reduces the capacity to sustain short-latency inhibition. This could be due to increased excitatory and/or reduced inhibitory transmission within reflex pathways (Robinson & Goldberger, 1985, 1986; De Leon *et al*., 1999; Tillakaratne *et al*., 2000, 2002; Ichiyama *et al*., 2011). However, studies in cats have found increased GABA synthesis in lumbar interneurons following complete thoracic SCI, consistent with increased inhibition (Tillakaratne *et al*., 2000, 2002). A recent study showed that the increased inhibitory phenotype at synaptic terminals of spinal interneurons, particularly in the dorsal and intermediate laminae, occurred through switching from excitatory to inhibitory after complete thoracic SCI in adult mice (Bertels *et al*., 2022). How these molecular synaptic changes influence reflex transmission is not known.

When considered as a whole, mid- and/or long-latency responses occurred less frequently after staggered hemisections in all four limbs (**Fig. 9**). Mid- and long-latency homonymous responses were significantly reduced after staggered hemisections in both hindlimbs, as well as mid-/long latency crossed responses in right hindlimb muscles but not left hindlimb muscles. When present, responses maintained their phase-dependent modulation. This suggests that pathways responsible for mid- and long-latency homonymous and crossed responses likely require supraspinal contributions/loops (Shimamura & Livingston, 1962; Shimamura & Aoki, 1969; Shimamura, 1973; Shimamura & Kogure, 1979; Shimamura *et al*., 1990). Supraspinal structures are known to modulate cutaneous reflexes and the stumbling corrective reaction during locomotion (Batson & Amassian, 1986; Fleshman *et al*., 1988; Amassian & Batson, 1988; Drew *et al*., 1996; Bretzner & Drew, 2005). Differences in crossed responses between left and right hindlimb muscles could reflect asymmetrical compensatory mechanisms within spinal sensorimotor circuits (Hultborn & Malmsten, 1983; Helgren & Goldberger, 1993; Frigon *et al*., 2009; Martinez *et al*., 2011; Audet *et al*., 2023). Homolateral and diagonal responses in forelimb muscles occurred less frequently in the intact state compared to homonymous and crossed responses and were strongly reduced after the first and second hemisections. Thus, mid- and long-latency responses in muscles of the four limbs are highly dependent on the integrity of spinal pathways and putative new connections are insufficient to support and restore them. The observation that phase-dependent modulation of reflex responses in fore- and hindlimb muscles was largely unaffected after staggered hemisections suggests that reflexes were modulated by spinal mechanisms, such as by CPGs and/or interactions with primary afferent inputs (Prochazka *et al*., 2002; Frigon & Rossignol, 2006; Frigon *et al*., 2021; Lalonde & Bui, 2021), although we cannot exclude a supraspinal contribution (Fleshman *et al*., 1988; Bretzner & Drew, 2005), particularly in forelimb muscles.

Lumbosacral circuits also undergo substantial reorganization following an incomplete thoracic SCI, such as after a lateral hemisection (Barriere *et al*., 2008; Frigon *et al*., 2009; Barrière *et al*., 2010; Martinez *et al*., 2011, 2012), or staggered lateral hemisections (Jane *et al*., 1964; Kato *et al*., 1984, 1985; Stelzner & Cullen, 1991; Courtine *et al*., 2008; Van Den Brand *et al*., 2012; Cowley *et al*., 2015; Audet *et al*., 2023) making them less dependent on supraspinal signals. At the circuit level, this can include collateral sprouting of primary afferent projections (Krenz & Weaver, 1998), which can affect reflex transmission and sensorimotor interactions between primary afferent inputs and CPGs. The mid- and long-latency reflex responses still observable in hindlimb muscles might reflect the reinforcement of remaining descending pathways, the activation of latent connections, and/or the establishment of new connections (Edgerton *et al*., 2004; Cai *et al*., 2006; Maier & Schwab, 2006; Basaldella *et al*., 2015; Higgin *et al*., 2020; Zavvarian *et al*., 2020).

Neural plasticity in lumbosacral circuits likely contributed to maintaining phase-dependent modulation of reflex responses in hindlimb muscles. The supraspinal contribution is more difficult to assess. Studies using staggered lateral hemisections have shown that transmission from the brainstem to lumbar levels remains possible (Cowley *et al*., 2008, 2010). Supraspinal pathways can synapse on propriospinal neurons undergoing reorganization above and/or below the lesion (Bareyre *et al*., 2004; Zaporozhets *et al*., 2006; Courtine *et al*., 2008; Cowley *et al*., 2008). New, short-distance propriospinal relays can allow bidirectional communication between cervical and lumbar levels (Courtine *et al*., 2008; Cowley *et al*., 2010; Zaporozhets *et al*., 2011; Laliberte *et al*., 2019). However, whether this transmission from supraspinal structures or from ascending somatosensory pathways through new propriospinal circuits can influence locomotion and reflex pathways in a meaningful way is not known. **Figure 11** schematically presents a scenario explaining changes in reflex responses to the four limbs after staggered hemisections with left SP nerve stimulation.

**Figure 11.**
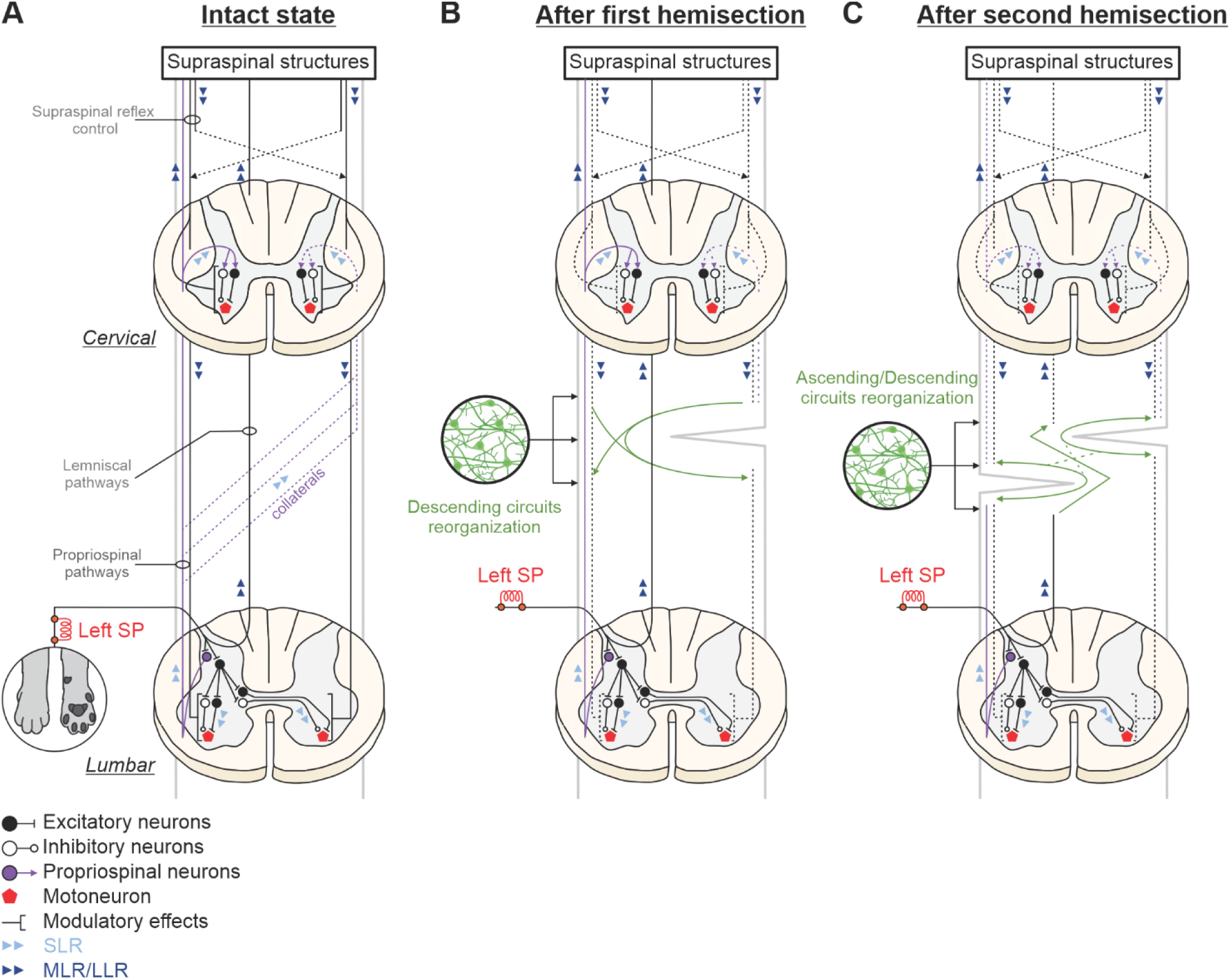
Schematic illustration of putative pathways and mechanisms contributing to cutaneous reflexes and their modulation before and after staggered hemisections. (**A**) In the intact state, afferents from the left superficial peroneal (SP) nerve contact spinal interneurons that project on motoneurons within the hemisegment (homonymous responses) at lumbar levels and commissural interneurons projecting contralaterally (crossed responses). SP nerve afferents also make contacts with propriospinal neurons that project to cervical levels, terminating ipsilaterally (homolateral responses) or contralaterally via collateral projections at different segments (diagonal responses). The pathways responsible for short-latency responses (SLRs) are mainly confined to spinal circuits, including SLRs in forelimb muscles. The pathways contributing to mid- and long-latency responses (MLRs/LLRs) transmit sensory information to supraspinal structures via long ascending projection neurons (propriospinal and/or dorsal lemniscal pathways) that project back to spinal circuits controlling the fore- and hindlimbs. (**B**) After the first hemisection (on the right side), SLRs in hindlimb muscles remain present although their response pattern can change due to functional changes in lumbosacral circuits. The occurrence of MLRs/LLRs decreases (dashed lines) due to disruptions in ascending and descending pathways to and from supraspinal structures. Spared supraspinal axons are potentially strengthened or sprout to form new connections to transmit descending signals. (**C**) After the second hemisection (on the left side), direct ascending and descending pathways are both disrupted, and reorganization of short propriospinal neurons is required to relay information through lesions, although their ability to do so is limited, leading to a considerable loss in MLRs and LLRs in all four limbs.

### Functional considerations

Stimulating the SP nerve elicits stumbling corrective and preventive reactions during the swing and stance phases of locomotion, respectively, as observed in both cats and humans (Forssberg *et al*., 1977; Prochazka *et al*., 1978; Forssberg, 1979; Duysens & Loeb, 1980; Wand *et al*., 1980; Schillings *et al*., 1996; Van Wezel *et al*., 1997; Zehr *et al*., 1997; Quevedo *et al*., 2005*b*, 2005*a*). Short-latency responses can rapidly modify limb trajectory and the locomotor pattern. During the swing phase, homonymous P1 responses in flexor muscles allow the stimulated limb to move the foot away and over a simulated obstacle. In contrast, during the stance phase, homonymous N1 responses reduce the activity of extensor muscles, briefly pausing forward progression and potentially lowering the center of gravity. After SCI, the ability to maintain balance and interlimb coordination during dynamic forward progression is altered or impaired, making adequate responses to perturbations even more essential (Ryu & Kuo, 2021). However, the ability to quickly respond to a perturbation is impaired after SCI. The latencies of short-latency responses in the ipsilesional right hindlimb slightly increased early after the first hemisection before returning toward intact values after eight weeks (**Fig. 10**). After an incomplete lateral SCI, cats shift their weight support to the contralesional hindlimb, which spends more time on the ground (Martinez *et al*., 2011, 2012; Audet *et al*., 2023). We observed shorter N1 response durations in extensors of the left contralesional hindlimb after the first hemisection (**Table 5**). This could be a compensatory mechanism to reinforce weight support. In a similar vein, Frigon and Rossignol (2008) showed the appearance of homonymous P1 responses in extensor muscles after spinal transection, instead of N1 responses.

While short-latency reflex responses can rapidly alter limb trajectory of the stimulated hindlimb, mid- and long-latency responses can be integrated into descending motor commands from the brain (Pruszynski & Scott, 2012), to assist in postural control for example. In the present study, the loss of mid- and longer-latency responses after staggered hemisections correlated with altered and/or impaired coordination of the fore- and hindlimbs and in dynamic balance. Studies in humans showed that cutaneous reflexes were modulated as a function of a real or perceived threat to stability (Llewellyn *et al*., 1990; Haridas *et al*., 2005, 2006, 2008; Lamont & Zehr, 2006, 2007). Thus, the loss of balance/stability can lead to reflex changes. We recently proposed in intact and spinal-transected cats that a signal related to an increase in left-right asymmetry, which affects walking stability (Dambreville *et al*., 2015; Huijben *et al*., 2018), reduced cutaneous reflex amplitudes during split-belt locomotion (Hurteau & Frigon, 2018; Mari *et al*., 2023). Single or staggered lateral hemisections induce temporal and spatial asymmetries in the locomotor pattern in cats (Kato *et al*., 1985; Martinez *et al*., 2011, 2013; Audet *et al*., 2023). Consequently, somatosensory signals related to these asymmetries in the locomotor pattern could have contributed to the loss/reduction of reflex responses after staggered hemisections.

### Limitations and concluding remarks

In conclusion, we showed that cutaneous reflex pathways from the foot dorsum projecting to the four limbs undergo considerable changes after staggered hemisections. These changes, especially a reduction/loss in their occurrence, correlated with impaired balance and fore-hind coordination. We are not proposing that changes in cutaneous reflexes led to these impairments but the changes in reflex pathways reflect the loss of communication between lumbosacral levels and circuits located at cervical and supraspinal levels. Many factors likely contributed to reflex changes, as discussed, but some are difficult to control experimentally. For example, cats have varying levels of physical activity after spinal lesions, with some more active than others. Before and after staggered hemisections, cats performed a variety of treadmill and walkway tasks to answer other scientific questions (Lecomte *et al*., 2022, 2023; Merlet *et al*., 2022; Audet *et al*., 2023; Mari *et al*., 2023). We believe that these tasks standardized the level of physical activity across animals, providing a minimal baseline. Lesion extent and the amount of secondary damage is also difficult to control, which undoubtedly affects locomotor recovery and changes in spinal pathways/circuits that contribute to reflexes and their modulation. We are currently investigating reflexes in the four limbs using the same dual lesion paradigm with stimulation of forelimb cutaneous afferents. Quantifying cutaneous reflex responses in the four limbs could be used as biomarkers to evaluate the efficacy of various therapeutic approaches (e.g., epidural stimulation of the spinal cord) in restoring transmission in ascending and descending spinal pathways and in sensorimotor control mechanisms.

## Support or grant information

This work was supported by a grant from the National Institutes of Health: R01 NS110550 to AF, IAR and BIP. AF is a Fonds de Recherche-Santé Quebec (FRQS) Senior Research Scholar. JA and JH were supported by FRQS doctoral scholarships and ANM by a FRQS postdoctoral scholarship.

## Author contributions

SM, IAR, BIP, and AF contributed to conception and design of the study. SM, CL, AM, JA, SY, OE, GG, CN and JH conducted the research. SM organized the database and performed the data and statistical analysis. SM and AF wrote the first draft of the manuscript. All authors contributed to manuscript revision, read, and approved the final version.

## Acknowledgments

We thank Philippe Drapeau for providing data acquisition and analysis software, developed in the Rossignol and Drew laboratories at the Université de Montréal. We thank the Biostatistics department of the Centre de Recherche du Centre Hospitalier Universitaire de Sherbrooke for statistical assisance.

## Data availability statement

The raw data supporting the conclusions of this article will be made available by the authors, without undue reservation.

## Ethics statement

The animal study was reviewed and approved by the Animal Care Committee of the Université de Sherbrooke.

## Competing statement

The authors declare no competing financial interests.

## REFERENCES

Amassian VE & Batson D (1988). Long loop participation of red nucleus in contact placing in the adult cat with facilitation by tactile input at the spinal level. Behav Brain Res 28, 225–232.

Audet J, Harnie J, Lecomte CG, Mari S, Merlet AN, Prilutsky BI, Rybak IA & Frigon A (2022). Control of Forelimb and Hindlimb Movements and Their Coordination during Quadrupedal Locomotion across Speeds in Adult Spinal Cats. J Neurotrauma 39, 1113–1131.

Audet J, Yassine S, Lecomte CG, Mari S, Soucy F, Morency C, Merlet AN, Harnie J, Beaulieu C, Gendron L, Rybak IA, Prilutsky BI & Frigon A (2023). Spinal sensorimotor circuits play a prominent role in hindlimb locomotor recovery after staggered thoracic lateral hemisections but cannot restore posture and interlimb coordination during quadrupedal locomotion in adult cats. eneuro 10, 0191– 23.

Barbeau H, Fung J, Leroux A & Ladouceur M (2002). A review of the adaptability and recovery of locomotion after spinal cord injury. In Progress in Brain Research, pp. 9–25. Elsevier Science.

Bareyre FM, Kerschensteiner M, Raineteau O, Mettenleiter TC, Weinmann O & Schwab ME (2004). The injured spinal cord spontaneously forms a new intraspinal circuit in adult rats. Nat Neurosci 7, 269– 277.

Barrière G, Frigon A, Leblond H, Provencher J & Rossignol S (2010). Dual Spinal Lesion Paradigm in the Cat: Evolution of the Kinematic Locomotor Pattern. J Neurophysiol 104, 1119–1133.

Barriere G, Leblond H, Provencher J & Rossignol S (2008). Prominent Role of the Spinal Central Pattern Generator in the Recovery of Locomotion after Partial Spinal Cord Injuries. J Neurosci 28, 3976– 3987.

Basaldella E, Takeoka A, Sigrist M & Arber S (2015). Multisensory Signaling Shapes Vestibulo-Motor Circuit Specificity. Cell 163, 301–312.

Batson DE & Amassian VE (1986). A dynamic role of rubral neurons in contact placing by the adult cat. J Neurophysiol 56, 835–856.

Berens P (2009). CircStat: A MATLAB Toolbox for Circular Statistics. J Stat Softw.

Bernard G, Bouyer L, Provencher J & Rossignol S (2007). Study of cutaneous reflex compensation during locomotion after nerve section in the cat. J Neurophysiol 97, 4173–4185.

Bertels H, Vicente-Ortiz G, El Kanbi K & Takeoka A (2022). Neurotransmitter phenotype switching by spinal excitatory interneurons regulates locomotor recovery after spinal cord injury. Nat Neurosci 25, 617–629.

Bouyer LJG & Rossignol S (2003*a*). Contribution of Cutaneous Inputs from the Hindpaw to the Control of Locomotion. I. Intact Cats. J Neurophysiol 90, 3625–3639.

Bouyer LJG & Rossignol S (2003*b*). Contribution of Cutaneous Inputs from the Hindpaw to the Control of Locomotion. II. Spinal Cats. J Neurophysiol 90, 3640–3653.

Bretzner F & Drew T (2005). Motor cortical modulation of cutaneous reflex responses in the hindlimb of the intact cat. J Neurophysiol 94, 673–687.

Budakova NN & Shik ML (1974). Stepping movements of the forelimbs and the Schiff-Sherrington phenomenon. Bull Exp Biol Med 77, 222–225.

Cai LL, Courtine G, Fong AJ, Burdick JW, Roy RR & Edgerton VR (2006). Plasticity of functional connectivity in the adult spinal cord. Philos Trans R Soc B Biol Sci 361, 1635–1646.

Calancie B, Molano MR & Broton JG (2002). Interlimb reflexes and synaptic plasticity become evident months after human spinal cord injury. Brain 125, 1150–1161.

Cartmill M, Lemelin P & Schmitt D (2002). Support polygons and symmetrical gaits in mammals. Zool J Linn Soc 136, 401–420.

Côté M-P, Detloff MR, Wade RE, Lemay MA & Houlé JD (2012). Plasticity in ascending long propriospinal and descending supraspinal pathways in chronic cervical spinal cord injured rats. Front Physiol 3, 00330.

Courtine G, Song B, Roy RR, Zhong H, Herrmann JE, Ao Y, Qi J, Edgerton VR & Sofroniew MV (2008). Recovery of supraspinal control of stepping via indirect propriospinal relay connections after spinal cord injury. Nat Med 14, 69–74.

Cowley KC, MacNeil BJ, Chopek JW, Sutherland S & Schmidt BJ (2015). Neurochemical excitation of thoracic propriospinal neurons improves hindlimb stepping in adult rats with spinal cord lesions. Exp Neurol 264, 174–187.

Cowley KC, Zaporozhets E & Schmidt BJ (2008). Propriospinal neurons are sufficient for bulbospinal transmission of the locomotor command signal in the neonatal rat spinal cord. J Physiol 586, 1623– 1635.

Cowley KC, Zaporozhets E & Schmidt BJ (2010). Propriospinal transmission of the locomotor command signal in the neonatal rat. Ann N Y Acad Sci 1198, 42–53.

Dambreville C, Labarre A, Thibaudier Y, Hurteau MF & Frigon A (2015). The spinal control of locomotion and step-to-step variability in left-right symmetry from slow to moderate speeds. J Neurophysiol 114, 1119–1128.

De Leon RD, Tamaki H, Hodgson JA, Roy RR & Edgerton VR (1999). Hindlimb Locomotor and Postural Training Modulates Glycinergic Inhibition in the Spinal Cord of the Adult Spinal Cat. J Neurophysiol 82, 359–369.

Deliagina TG, Beloozerova IN, Orlovsky GN & Zelenin PV (2014). Contribution of supraspinal systems to generation of automatic postural responses. Front Integr Neurosci 8, 00076.

Drew T, Cabana T & Rossignol S (1996). Responses of medullary reticulospinal neurones to stimulation of cutaneous limb nerves during locomotion in intact cats. Exp Brain Res 111, 153–168.

Duysens J & Loeb GE (1980). Modulation of ipsi- and contralateral reflex responses in unrestrained walking cats. J Neurophysiol 44, 1024–1037.

Duysens J & Stein RB (1978). Reflexes induced by nerve stimulation in walking cats with implanted cuff electrodes. Exp Brain Res 32, 213–224.

Edgerton VR, Tillakaratne NJK, Bigbee AJ, De Leon RD & Roy RR (2004). Plasticity of the spinal neural circuitry after injury*. Annu Rev Neurosci 27, 145–167.

English AW (1979). Interlimb Coordination During Stepping in the Cat: an Electromyographic Analysis. J Neurophysiol 42, 229–243.

English AW (1980). Interlimb coordination during stepping in the cat: effects of dorsal column section. J Neurophysiol 44, 270–279.

English AW (1985). Interlimb coordination during stepping in the cat: The role of the dorsal spinocerebellar tract. Exp Neurol 87, 96–108.

English AW & Lennard PR (1982). Interlimb Coordination During Stepping in the Cat : In-Phase Stepping and Gait Transitions. Brain Res 245, 353–364.

Fleshman JW, Rudomin P & Burke RE (1988). Supraspinal control of a short-latency cutaneous pathway to hindlimb motoneurons. Exp Brain Res 69, 449–459.

Forssberg H (1979). Stumbling Corrective Reaction: A Phase-Dependent Compensatory Reaction During Locomotion. J Neurophysiol 42, 36–953.

Forssberg H, Grillner S & Rossignol S (1977). Phasic gain control of reflexes from the dorsum of the paw during spinal locomotion. Brain Res 132, 121–139.

Fouad K & Pearson K (2004). Restoring walking after spinal cord injury. Prog Neurobiol 73, 107–126.

Frigon A (2011). Interindividual variability and its implications for locomotor adaptation following peripheral nerve and/or spinal cord injury. Prog Brain Res 188, 101–118.

Frigon A (2017). The neural control of interlimb coordination during mammalian locomotion. J Neurophysiol 117, 2224–2241.

Frigon A, Akay T & Prilutsky BI (2021). Control of Mammalian Locomotion by Somatosensory Feedback. Compr Physiol 12, 2877–2947.

Frigon A, Barrière G, Leblond H & Rossignol S (2009). Asymmetric Changes in Cutaneous Reflexes After a Partial Spinal Lesion and Retention Following Spinalization During Locomotion in the Cat. J Neurophysiol 102, 2667–2680.

Frigon A & Rossignol S (2006). Functional plasticity following spinal cord lesions. Prog Brain Res 157, 231– 260.

Frigon A & Rossignol S (2007). Short-Latency Crossed Inhibitory Responses in Extensor Muscles During Locomotion in the Cat. J Neurophysiol 99, 989–998.

Frigon A & Rossignol S (2008). Adaptive changes of the locomotor pattern and cutaneous reflexes during locomotion studied in the same cats before and after spinalization. J Physiol 586, 2927–2945.

Frigon A & Rossignol S (2009). Partial denervation of ankle extensors prior to spinalization in cats impacts the expression of locomotion and the phasic modulation of reflexes. Neuroscience 158, 1675–1690.

Frigon A, Thibaudier Y, Hurteau MF, Telonio A, Dambreville C & Kuczynski V (2014). The control of interlimb coordination during left-right and transverse split-belt locomotion in intact and spinal cord-injured cats. Biosyst Biorobotics 7, 29–34.

Fung J & Macpherson JM (1999). Attributes of Quiet Stance in the Chronic Spinal Cat. J Neurophysiol 82, 3056–3065.

Fuwa T, Shimamura M & Tanaka I (1991). Analysis of the forelimb crossed extension reflex in thalamic cats during stepping. Neurosci Res 9, 257–269.

Goldberger ME (1977). Locomotor recovery after unilateral hindlimb deafferentation in cats. Brain Res 123, 59–74.

Górska T, Chojnicka-Gittins B, Majczyński H & Zmysłowski W (2013). Changes in forelimb–hindlimb coordination after partial spinal lesions of different extent in the rat. Behav Brain Res 239, 121–138.

Gossard JP, Delivet-Mongrain H, Martinez M, Kundu A, Escalona M & Rossignol S (2015). Plastic changes in lumbar locomotor networks after a partial spinal cord injury in cats. J Neurosci 35, 9446–9455.

Haridas C & Zehr EP (2003). Coordinated Interlimb Compensatory Responses to Electrical Stimulation of Cutaneous Nerves in the Hand and Foot during Walking. J Neurophysiol 90, 2850–2861.

Haridas C, Zehr EP & Misiaszek JE (2005). Postural uncertainty leads to dynamic control of cutaneous reflexes from the foot during human walking. Brain Res 1062, 48–62.

Haridas C, Zehr EP & Misiaszek JE (2006). Context-dependent modulation of interlimb cutaneous reflexes in arm muscles as a function of stability threat during walking. J Neurophysiol 96, 3096–3103.

Haridas C, Zehr EP & Misiaszek JE (2008). Adaptation of cutaneous stumble correction when tripping is part of the locomotor environment. J Neurophysiol 99, 2789–2797.

Helgren ME & Goldberger ME (1993). The recovery of postural reflexes and locomotion following low thoracic hemisection in adult cats involves compensation by undamaged primary afferent pathways. Exp Neurol 123, 17–34.

Higgin D, Krupka A, Maghsoudi OH, Klishko AN, Nichols TR, Lyle MA, Prilutsky BI & Lemay MA (2020). Adaptation to slope in locomotor-trained spinal cats with intact and self-reinnervated lateral gastrocnemius and soleus muscles. J Neurophysiol 123, 70–89.

Huijben B, Van Schooten KS, Van Dieën JH & Pijnappels M (2018). The effect of walking speed on quality of gait in older adults. Gait Posture 65, 112–116.

Hultborn H & Malmsten J (1983). Changes in segmental reflexes following chronic spinal cord hemisection in the cat: Acta Physiol Scand 119, 405–422.

Hurteau M & Frigon A (2018). A spinal mechanism related to left–right symmetry reduces cutaneous reflex modulation independently of speed during split-belt locomotion. J Neurosci 38, 10314–10328.

Hurteau M, Thibaudier Y, Dambreville C, Danner SM, Rybak IA & Frigon A (2018). Intralimb and Interlimb Cutaneous Reflexes during Locomotion in the Intact Cat. J Neurosci 38, 4104–4122.

Hurteau M-F, Thibaudier Y, Dambreville C, Chraibi A, Desrochers E, Telonio A & Frigon A (2017). Nonlinear Modulation of Cutaneous Reflexes with Increasing Speed of Locomotion in Spinal Cats. J Neurosci 37, 3896–3912.

Ichiyama RM, Broman J, Roy RR, Zhong H, Edgerton VR & Havton LA (2011). Locomotor Training Maintains Normal Inhibitory Influence on Both Alpha- and Gamma-Motoneurons after Neonatal Spinal Cord Transection. J Neurosci 31, 26–33.

Jane JA, Evans JP & Fisher LE (1964). An Investigation Concerning the Restitution of Motor Function Following Injury to the Spinal Cord. J Neurosurg 21, 167–171.

Kato M, Murakami S, Hirayama H & Hikino K (1985). Recovery of postural control following chronic bilateral hemisections at different spinal cord levels in adult cats. Exp Neurol 90, 350–364.

Kato M, Murakami S, Yasuda K & Hirayama H (1984). Disruption of fore- and hindlimb coordination during overground locomotion in cats with bilateral serial hemisection of the spinal cord. Neurosci Res 2, 27–47.

Kloos A, Fisher L, Detloff M, Hassenzahl D & Basso D (2005). Stepwise motor and all-or-none sensory recovery is associated with nonlinear sparing after incremental spinal cord injury in rats. Exp Neurol 191, 251–265.

Krenz NR & Weaver LC (1998). Sprouting of primary afferent fibers after spinal cord transection in the rat. Neuroscience 85, 443–458.

LaBella LA, Niechaj A & Rossignol S (1992). Low-threshold, short-latency cutaneous reflexes during fictive locomotion in the “semi-chronic” spinal cat. Exp Brain Res 91, 236–248.

Laliberte AM, Goltash S, Lalonde NR & Bui TV (2019). Propriospinal Neurons: Essential Elements of Locomotor Control in the Intact and Possibly the Injured Spinal Cord. Front Cell Neurosci 13, 00512.

Lalonde NR & Bui TV (2021). Do spinal circuits still require gating of sensory information by presynaptic inhibition after spinal cord injury? Curr Opin Physiol 19, 113–118.

Lamont EV & Zehr EP (2006). Task-specific modulation of cutaneous reflexes expressed at functionally relevant gait cycle phases during level and incline walking and stair climbing. Exp Brain Res 173, 185–192.

Lamont EV & Zehr EP (2007). Earth-referenced handrail contact facilitates interlimb cutaneous reflexes during locomotion. J Neurophysiol 98, 433–442.

Lecomte CG, Audet J, Harnie J & Frigon A (2021). A Validation of Supervised Deep Learning for Gait Analysis in the Cat. Front Neuroinformatics 15, 712623.

Lecomte CG, Mari S, Audet J, Merlet AN, Harnie J, Beaulieu C, Abdallah K, Gendron L, Rybak IA, Prilutsky BI & Frigon A (2022). Modulation of the gait pattern during split-belt locomotion after lateral spinal cord hemisection in adult cats. J Neurophysiol 128, 1593–1616.

Lecomte CG, Mari S, Audet J, Yassine S, Merlet AN, Morency C, Harnie J, Beaulieu C, Gendron L & Frigon A (2023). Neuromechanical Strategies for Obstacle Negotiation during Overground Locomotion following Incomplete Spinal Cord Injury in Adult Cats. J Neurosci 43, 5623–5641.

Llewellyn M, Yang JF & Prochazka A (1990). Human H-reflexes are smaller in difficult beam walking than in normal treadmill walking. Exp Brain Res 83, 22–28.

Loeb GE (1993). The distal hindlimb musculature of the cat: interanimal variability of locomotor activity and cutaneous reflexes. Exp Brain Res 96, 125–140.

Macpherson JM & Fung J (1999). Weight Support and Balance During Perturbed Stance in the Chronic Spinal Cat. J Neurophysiol 82, 3066–3081.

Maier IC & Schwab ME (2006). Sprouting, regeneration and circuit formation in the injured spinal cord: factors and activity. Philos Trans R Soc B Biol Sci 361, 1611–1634.

Mari S, Lecomte CG, Merlet AN, Audet J, Harnie J, Rybak IA, Prilutsky BI & Frigon A (2023). A sensory signal related to left-right symmetry modulates intra- and interlimb cutaneous reflexes during locomotion in intact cats. Front Syst Neurosci 17, 1199079.

Martinez M, Delivet-Mongrain H, Leblond H & Rossignol S (2011). Recovery of hindlimb locomotion after incomplete spinal cord injury in the cat involves spontaneous compensatory changes within the spinal locomotor circuitry. J Neurophysiol 106, 1969–1984.

Martinez M, Delivet-Mongrain H, Leblond H & Rossignol S (2012). Incomplete spinal cord injury promotes durable functional changes within the spinal locomotor circuitry. J Neurophysiol 108, 124–134.

Martinez M, Delivet-Mongrain H & Rossignol S (2013). Treadmill training promotes spinal changes leading to locomotor recovery after partial spinal cord injury in cats. J Neurophysiol 109, 2909–2922.

Mathis A, Mamidanna P, Cury KM, Abe T, Murthy VN, Mathis MW & Bethge M (2018). DeepLabCut: markerless pose estimation of user-defined body parts with deep learning. Nat Neurosci 21, 1281– 1289.

Matthews PBC (1986). Observations of the automatic compensation of reflex gain on varying the pre- existing level on motor discharge in man. J Physiol 374, 73–90.

Merlet AN, Harnie J & Frigon A (2021). Inhibition and facilitation of the spinal locomotor central pattern generator and reflex circuits by somatosensory feedback from the lumbar and perineal regions after spinal cord injury. Front Neurosci 15, 1–14.

Merlet AN, Harnie J, Macovei M, Doelman A, Gaudreault N & Frigon A (2020). Mechanically stimulating the lumbar region inhibits locomotor-like activity and increases the gain of cutaneous reflexes from the paws in spinal cats. J Neurophysiol 123, 1026–1041.

Merlet AN, Jéhannin P, Mari S, Lecomte CG, Audet J, Harnie J, Rybak IA, Prilutsky BI & Frigon A (2022). Sensory perturbations from hindlimb cutaneous afferents generate coordinated functional responses in all four limbs during locomotion in intact cats. eNeuro 9, 0178–22.

Muir GD & Steeves JD (1995). Phasic cutaneous input facilitates locomotor recovery after incomplete spinal injury in the chick. J Neurophysiol 74, 358–368.

Murray KC, Nakae A, Stephens MJ, Rank M, D’Amico J, Harvey PJ, Li X, Harris RLW, Ballou EW, Anelli R, Heckman CJ, Mashimo T, Vavrek R, Sanelli L, Gorassini MA, Bennett DJ & Fouad K (2010). Recovery of motoneuron and locomotor function after spinal cord injury depends on constitutive activity in 5-HT2C receptors. Nat Med 16, 694–700.

Orsal D, Cabelguen JM & Perret C (1990). Interlimb coordination during fictive locomotion in the thalamic cat. Exp Brain Res 82, 536–546.

Percie Du Sert N et al. (2020). The ARRIVE guidelines 2.0: Updated guidelines for reporting animal research*. J Cereb Blood Flow Metab 40, 1769–1777.

Pratt CA, Chanaud CM & Loeb GE (1991). Functionally complex muscles of the cat hindlimb IV. Intramuscular distribution of movement command signals and cutaneous reflexes in broad, bifunctional thigh muscles. Exp Brain Res 85, 281–299.

Pratt CA, Fung J & Macpherson JM (1994). Stance control in the chronic spinal cat. J Neurophysiol 71, 1981– 1985.

Prochazka A, Mushahwar V & Yakovenko S (2002). Activation and coordination of spinal motoneuron pools after spinal cord injury. Prog Brain Res 137, 109–124.

Prochazka A, Sontag K-H & Wand P (1978). Motor reactions to perturbations of gait : proprioceptive and somesthetic involvement. Neurosci Lett 7, 35–39.

Pruszynski JA & Scott SH (2012). Optimal feedback control and the long-latency stretch response. Exp Brain Res 218, 341–359.

Quevedo J, Stecina K, Gosgnach S & McCrea DA (2005*a*). Stumbling corrective reaction during fictive locomotion in the cat. J Neurophysiol 94, 2045–2052.

Quevedo J, Stecina K & McCrea DA (2005*b*). Intracellular analysis of reflex pathways underlying the stumbling corrective reaction during fictive locomotion in the cat. J Neurophysiol 94, 2053–2062.

Robinson GA & Goldberger ME (1985). Interfering with inhibition may improve motor function. Brain Res 346, 400–403.

Robinson GA & Goldberger ME (1986). The development and recovery of motor function in spinal cats. Exp Brain Res 62, 387–400.

Rossignol S, Dubuc R & Gossard JP (2006). Dynamic sensorimotor interactions in locomotion. Physiol Rev 86, 89–154.

Rossignol S & Frigon A (2011). Recovery of locomotion after spinal cord injury: Some facts and mechanisms. Annu Rev Neurosci 34, 413–440.

Ryu HX & Kuo AD (2021). An optimality principle for locomotor central pattern generators. Sci Rep 11, 1–18.

Schillings AM, Van Wezel BMH & Duysens J (1996). Mechanically induced stumbling during human treadmill walking. J Neurosci Methods 67, 11–17.

Shimamura M (1973). Spino-bulbo-spinal and propriospinal reflexes in various vertebrates. Brain Res 64, 141–165.

Shimamura M & Aoki M (1969). Effects of spino-bulbo-spinal reflex volleys on flexor motoneurons of hindlimb in the cat. Brain Res 16, 333–349.

Shimamura M & Kogure I (1979). Reticulospinal tracts involved in the spino-bulbo-spinal reflex in cats. Brain Res 172, 13–21.

Shimamura M & Livingston R (1962). Longitudinal conduction systems serving spinal and brainstem coordination-spino-bulbo-spinal reflex. J Physiol Soc Jpn 33, 225–233.

Shimamura M, Tanaka I & Fuwa T (1990). Comparison between spino-bulbo-spinal and propriospinal reflexes in thalamic cats during stepping. Neurosci Res 7, 358–368.

Smith RR, Shum-Siu A, Baltzley R, Bunger M, Baldini A, Burke DA & Magnuson DSK (2006). Effects of Swimming on Functional Recovery after Incomplete Spinal Cord Injury in Rats. J Neurotrauma 23, 908–919.

Stelzner DJ & Cullen JM (1991). Do Propriospinal Projections Contribute to Hindlimb Recovery when All Long Tracts Are Cut in Neonatal or Weanling Rats? Exp Neurol 114, 193–205.

Takeoka A, Vollenweider I, Courtine G & Arber S (2014). Muscle Spindle Feedback Directs Locomotor Recovery and Circuit Reorganization after Spinal Cord Injury. Cell 159, 1626–1639.

Thibaudier Y & Frigon A (2014). Spatiotemporal control of interlimb coordination during transverse split-belt locomotion with 1:1 or 2:1 coupling patterns in intact adult cats. J Neurophysiol 112, 2006– 2018.

Thibaudier Y, Hurteau MF, Dambreville C, Chraibi A, Goetz L & Frigon A (2017). Interlimb Coordination during Tied-Belt and Transverse Split-Belt Locomotion before and after an Incomplete Spinal Cord Injury. J Neurotrauma 34, 1751–1765.

Tillakaratne NJK, De Leon RD, Hoang TX, Roy RR, Edgerton VR & Tobin AJ (2002). Use-Dependent Modulation of Inhibitory Capacity in the Feline Lumbar Spinal Cord. J Neurosci 22, 3130–3143.

Tillakaratne NJK, Mouria M, Ziv NB, Roy RR, Edgerton VR & Tobin AJ (2000). Increased expression of glutamate decarboxylase (GAD67) in feline lumbar spinal cord after complete thoracic spinal cord transection. J Neurosci Res 60, 219–230.

Van Den Brand R, Heutschi J, Barraud Q, DiGiovanna J, Bartholdi K, Huerlimann M, Friedli L, Vollenweider I, Moraud EM, Duis S, Dominici N, Micera S, Musienko P & Courtine G (2012). Restoring voluntary control of locomotion after paralyzing spinal cord injury. Science 336, 1182–1185.

Van Hedel HJA & Dietz V (2010). Rehabilitation of locomotion after spinal cord injury. Restor Neurol Neurosci 28, 123–134.

Van Wezel BMH, Ottenhoff FAM & Duysens J (1997). Dynamic Control of Location-Specific Information in Tactile Cutaneous Reflexes from the Foot during Human Walking. J Neurosci 17, 3804–3814.

Wand P, Prochazka A & Sontag K-H (1980). Neuromuscular Responses to Gait Perturbations in Freely Moving Cats*. Exp Brain Res 38, 109–114.

Zaporozhets E, Cowley KC & Schmidt BJ (2006). Propriospinal neurons contribute to bulbospinal transmission of the locomotor command signal in the neonatal rat spinal cord. J Physiol 572, 443– 458.

Zaporozhets E, Cowley KC & Schmidt BJ (2011). Neurochemical excitation of propriospinal neurons facilitates locomotor command signal transmission in the lesioned spinal cord. J Neurophysiol 105, 2818–2829.

Zavvarian M-M, Hong J & Fehlings MG (2020). The Functional Role of Spinal Interneurons Following Traumatic Spinal Cord Injury. Front Cell Neurosci 14, 00127.

Zehr EP, Komiyama T & Stein RB (1997). Cutaneous reflexes during human gait: Electromyographic and kinematic responses to electrical stimulation. J Neurophysiol 77, 3311–3325.

